# TRiPPing the sensors: The osmosensing pathway of Polycystin 2

**DOI:** 10.1101/2023.05.09.540007

**Authors:** K.M. Márquez-Nogueras, R.M. Knutila, V. Vuchkosvka, IY. Kuo

**Author notes:** Corresponding Author: Ivana Y. Kuo.

## Abstract

Mutations to polycystin-2 (PC2), a non-selective cation permeant transient receptor potential channel, results in polycystic kidney disease (PKD). Despite the disease relevance of PC2, the physiological agonist that activates PC2 has remained elusive. As one of the earliest symptoms in PKD is a urine concentrating deficiency, we hypothesized that shifts in osmolarity experienced by the collecting duct cells would activate PC2 and loss of PC2 would prevent osmosensing. We found that mice with inducible PC2 knocked out (KO) in renal tubules had dilute urine. Hyperosmotic stimuli induced a rise in endoplasmic reticulum (ER)-mediated cytosolic calcium which was absent in PC2 KO mice and PC2 KO cells. A pathologic point mutation that prevents ion flux through PC2 inhibited the calcium rise, pointing to the centrality of PC2 in the osmotic response. To understand how an extracellular stimulus activated ER-localized PC2, we examined microtubule-ER dynamics, and found that the osmotically induced calcium increase was preceded by microtubule destabilization. This was due to a novel interaction between PC2 and the microtubule binding protein MAP4 that tethers the microtubules to the ER. Finally, disruption of the MAP4-PC2 interaction prevented incorporation of the water channel aquaporin 2 following a hyperosmotic challenge, in part explaining the dilute urine. Our results demonstrate that MAP4-dependent microtubule stabilization of ER-resident PC2 is required for PC2 to participate in the osmosensing pathway. Moreover, osmolarity represents a *bona fide* physiological stimulus for ER-localized PC2 and loss of PC2 in renal epithelial cells impairs osmosensing ability and urine concentrating capacity.

**Graphical Abstract:** 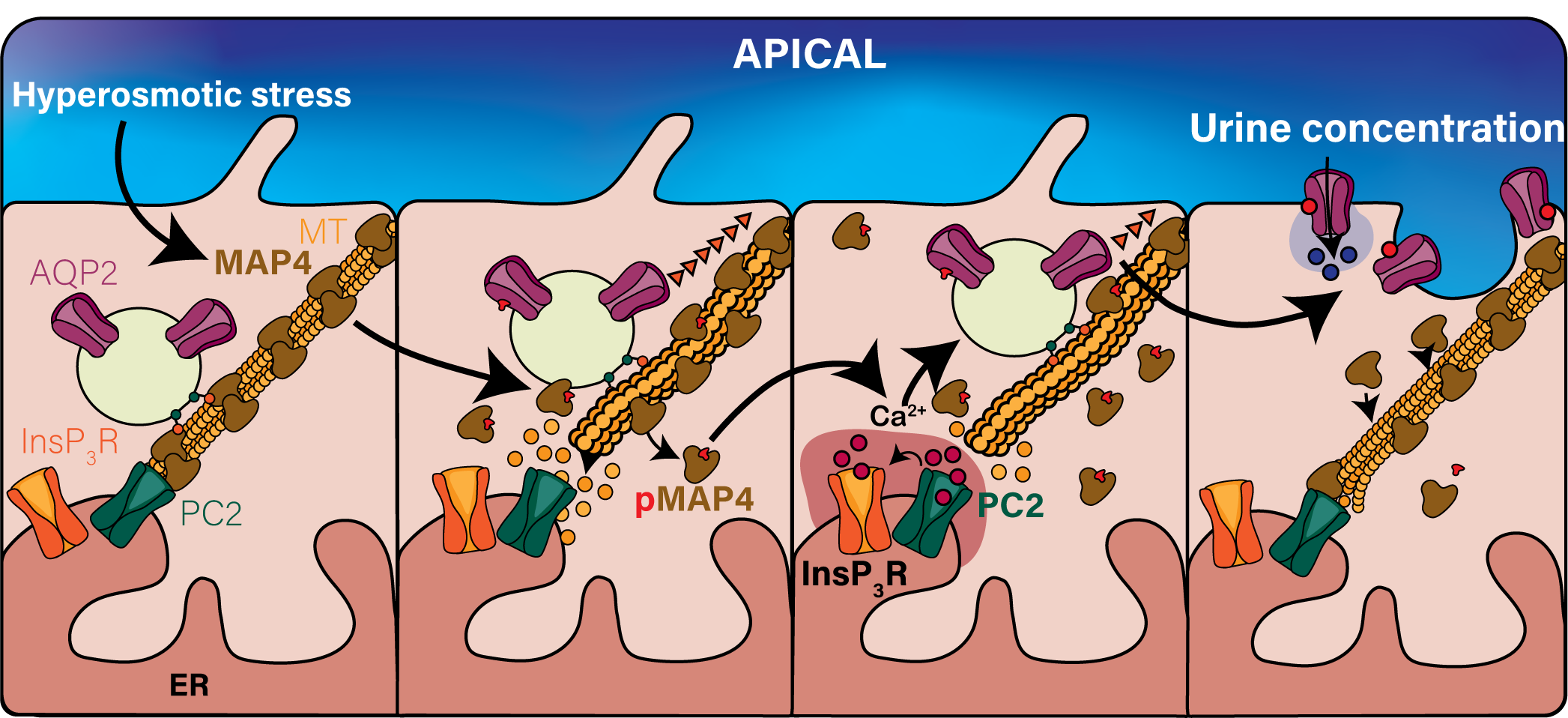

## Introduction

Mechano-transduction pathways couple extracellular cues, such as fluid flow and stretch to a signaling response that enable a cell to adequately respond to changes in the environment. These extracellular cues can be transmitted via ion channels on the plasma membrane such as piezo or transient receptor potential (TRP) channels [1, 2]. One such environment where extracellular cues must be rapidly converted into intracellular signals is the kidney, where urine is concentrated. The concentration of urine requires the selective salt and water permeabilities of the renal epithelial cells within the tubule. The active pumping of salts combined with the urea recycling help to generate the osmotic gradient that can reach up to ∼1,200 mOsm in the depth of the medulla in humans [3]. However, the osmolarity of the tubular fluid (i.e. urine) flowing through the renal tubules varies depending on a body’s demands. The final osmolarity of the urine as it exits the collecting duct varies between ∼50 to 1,200 mOsm [3]. Thus, medullary collecting duct cells need to quickly adapt to changes in tubular fluid osmolarity to maintain the medullary gradient. Disruption in the ability of the cell to adapt to these responses impair the kidney’s ability to concentrate urine [4].

A lack of urine concentrating ability is observed in many renal disorders including Autosomal Dominant Polycystic Kidney Disease (ADPKD), the leading genetic cause of renal failure which has no cure [5]. Although ADPKD is characterized by the formation of renal cysts throughout the nephron, including in the collecting duct, impaired urine concentration is an early symptom of the disease [6]. As urine flows, water is reabsorbed in the collecting duct which is partially mediated by aquaporin 2 (AQP2) which allows for water flux into the cell across the apical membrane [3, 7]. Expression and trafficking of AQP2 is canonically activated by arginine vasopressin (AVP) [8]. AVP binds to the vasopressin receptor type 2 (V2R) on the basolateral side of the membrane leading to increased cAMP inducing translation and vesicular trafficking of AQP2 [8]. Despite the physiological importance of urine concentration, the mechanosensitive signaling pathways that allow for collecting duct epithelial cells to respond to changes in urine osmolarity and how these pathways affect AQP2 insertion into the membrane in ADPKD models is not fully understood.

Polycystin 2 (PC2) is one of the two main genes which when mutated results in ADPKD [9]. PC2 is a non-selective cation channel that belongs to the TRP polycystin channel family and can reside on the primary cilia, plasma membrane and endoplasmic reticulum (ER) [10]. In the ER, PC2 can mediate calcium release either directly or via interactions with other calcium release channels [11–13]. Electrophysiological approaches have demonstrated that PC2 can mediate flux of sodium, potassium, and calcium across the ciliary or plasma membrane [14–18]. In primary cilia, PC2, either by itself or in complex with its interacting partner polycystin 1 (PC1), can act as a mechanosensitive channel that opens in response to changes to fluid flow [19]. Recent work demonstrated that in nodal cilia, PC2 is activated upon multiple bending of the cilia and is required for left-right patterning [20, 21]. However, a high number of ciliary bends are required to elicit the calcium signal. As there is no consensus for the agonist for PC2 in renal epithelial cells, it is unclear whether the bending experienced by the primary cilia under fluid flow of urine is the true physiologic signal required to activate a calcium signal from the PC2 complex.

As the majority of PC2 resides in the ER, we examined the contribution of PC2 to calcium signaling responses arising from the primary cilia, the plasma membrane, and the ER upon osmotic changes. We hypothesized that shifts in extracellular tonicity would act as a natural agonist for ER-localized PC2 in renal epithelial cells and that loss of PC2, as would be expected in the setting of ADPKD, would impair urine concentrating ability. Compared to the control (CTL) mice, we found that deletion of PC2 in the tubules of pre-cystic mice resulted in dilute urine. Hyperosmotic stimuli induced a calcium response in isolated kidney tubules and in a variety of cell lines. This response was absent in kidney tubules from pre-cystic PC2 KO mice and in cultured cell lines. Re-expression of full-length PC2 but not a pathogenic variant, restored the cytosolic calcium response. The calcium response was found to originate in the ER and not the primary cilia or plasma membrane. Mechanistically, we identified a novel molecular mechanism by which the osmotic stimuli activated PC2 calcium release. PC2 associated with the microtubule binding protein MAP4, which upon hyperosmotic stimuli, dissociated from the microtubule allowing for PC2 calcium release. The biological necessity of the interaction between PC2 and MAP4 to transduce osmotic stimuli was shown as trafficking of the AQP2 was abolished in the PC2 KO cells, which ultimately contributes to diminished urine concentration.

## Materials and methods

### Animal model

*Pkd2* floxed mice [22] (gift from Dr. Stefan Somlo, Yale University) were crossed with TetO-Cre and Pax8-rtTA mice (Jackson laboratories strain #006234 and #007176) to generate *Pkd2^FF^*TetO-Cre-Pax8 mice and TetO-Cre-Pax8 control mice. Some mice were further crossed with LSL-Salsa6f mice (Jackson laboratories strain #031968) to generate mice that expressed gCaMP6F and tdTomato upon Cre induction. At 6 weeks of age, mice were fed doxycycline chow (625 mg/kg) for 7 days to induce Cre expression. Male and female mice, 13-20 weeks of age (1.5-3 months post induction) were used in the subsequent experiments. All animal studies performed were done under approved Institutional Animal Care and Use Committee (IACUC) protocols at Loyola University Chicago.

### Urine analysis, metabolic cages

Animals provided with *ad lib* water were moved to a hydrophobic surface and urine collected in the morning within the first 3 min of the animal being removed from the cage. Urine volumes were between 50 to 100 μL. Urine was measured with a vapor osmometer (Wescor 5520 Vapor Pressure Osmometer). A subset of animals had body measurements conducted by NMR spectroscopy followed by metabolic cage analysis (TSE Systems) to measure energetic expenditure, food, and water intake over 5 days.

### Imaging of kidney sections

Animals were sedated with isoflurane and perfused with saline via the left ventricle. The kidneys were rapidly excised and dissected into ∼1mm coronal sections in ice cold PBS and then transferred to kidney tubule solution (mM: 120 NaCl; 3 KCl; 2 CaCl_2_; 2 KH_2_PO_4_ 2; 5 Glucose; 10 HEPES; pH 7.3). Sections were either incubated in Fluo4 (5 μM) for 30 min at room temperature (RT) or incubated without Fluo4-AM in gCaMP expressing kidneys. Sections were washed 2 times with fresh kidney tubule solution. Medullary tubules were identified by visual inspection based on anatomical landmarks. Images were acquired by incubating the tubules at 250 mOsm (5 min) followed by 400 mOsm stimuli (10 min). To ensure distal and collecting duct tubules were being assessed, only tubules that responded to exogenously applied vasopressin were included in the analysis. Ionomycin was added at the completion of the experiment.

### Cell culture and maintenance

Murine C2C12 myoblasts and Human Embryonic Kidney (HEK293T) cells were purchased from ATCC. HEK293 cells that contain all the isoforms of the InsP_3_R knocked out (known as 3KO-HEK cells) were a gift from Dr. David Yule from the University of Rochester [23]. Immortalized murine collecting duct cells (imCD3) cells were a gift from Indra Chandrasekar from Sanford Health. C2C12 and HEK293T cells were cultured in Dulbecco’s modified Eagle’s medium (DMEM) while imCD3 cells were cultured in F-12 media. All media was supplemented with 10% fetal bovine essence (FBE) and antibiotics at 37°C and incubated in a humidified atmosphere with 5% CO_2_ and 95% air. Cell cultures were kept to passages no higher than P12.

### Generation of PC2 and MAP-4 CRISPR KO cell lines

We tested two different CRISPR/Cas9 knockout All-In-One ZsGreen pClip lentivirus plasmids each containing a separate guide sequence directed at different *Pkd*2 and *MAP4* loci, along with a control template (Transomic Technologies). The lentiviruses were made by co-transfecting pRSV, pMDLg and pMD2.G along with pClip into HEK293T cells. The supernatant containing virus particles was harvested and used to transduce C2C12, imCD3, or HEK293T cells. Following 48 hours of transduction, ZsGreen fluorescent C2C12, imCD3 or HEK293T cells were sorted by flow cytometry and *Pkd2* single cell clones expanded. Selection of MAP4 from the mixed population was obtained by supplementing the media with 10 μM of blastocidin. Following expansion, the different cell lines were validated by western blot, qPCR and immunofluorescent assay.

### Measurement of intracellular calcium

We measured changes in calcium in the following cellular compartments: cytosol, ER, plasma membrane and cilia. Calcium measurements were measured using the following plasmids: gCAMP6F (cytosolic calcium), R-cepia (ER calcium, gift of Dr. Aleskey Zima) [24], CAAX-gCaMP7s (plasma membrane, gift of Dr. Jordan Beach) and Arl13B-gCaMP6F (cilia, gift of Dr. Aldebaran Hofer and made by the Yubin Zhao’s lab). Cells were plates on glass coverslips 48 hours prior to imaging. Cells were transiently transfected 24 hours after plating with 2 μg of the DNA of interest and 25 μL of PEI (concentration 1μg/μl stock) diluted in OPTI-MEM and incubated overnight. Regions of interest were drawn on individual cells to quantify calcium measurements in at least 3 independent biological replicates. Cells were perfused with a Warner gravity exchange fluid system (Warner instruments) for 2 min at 3ml/mn with 300 mOsm solution (mM: 130 NaCl; 2 CaCl_2_; 1 MgCl; 2 K_2_PO_4_ 2; 5 Glucose; 10 HEPES; 0.1 EGTA; pH 7.3) prior to imaging. Cells were then perfused with the specified osmolarity. Increase of osmolarity was done by increasing the concentration of NaCl or mannitol as described in results. The osmolarity of solutions was measured by a vapor osmometer (Wescor 5520 Vapor Pressure Osmometer). gCaMP6F was excited with a 488nM LED (Lumnecor Spectra X Lamp) and emitted fluorescence filtered with a band pass filter (515-530, Chrom. Images were acquired with a sCMOS camera (Orca Flash, Hamamatsu) on a wide-field fluorescence Zeiss microscope. Images were acquired at 15ms for a total of 3-6 minutes at RT.

### Western blot analysis

Total protein extracts were collected after incubating cells under the different osmotic conditions during 1 hour at 37°C and prepared by lysing cells with RIPA buffer (in mM: 10 Tris-Cl, 1 EDTA, 0.5 EGTA, 1% Triton-X, 0.1% sodium deoxycholate, 0.1% SDS, 140 NaCl) containing protease inhibitor cocktail (Sigma-Aldrich), and phosphatase inhibitors NaF and sodium orthovanadate (Alfa Aesar). Protein concentrations of the resulting supernatants were measured using the Pierce BCA Protein Assay Kit (Thermo Scientific). Equal amounts of protein (15-20 μg) were separated by SDS-PAGE (Bio-Rad, 4-20% gradient gels) and transferred to PVDF membranes via wet transfer. Membranes were probed overnight with the following primary antibodies: ⍺-tubulin (1:1,000, #2125S; Cell Signaling Technology), GAPDH (1:1,500, #6004-Ig; ProteinTech), PC2 (D-3, 1:500, sc-28331; Santa Cruz Biotechnology), MAP4 (1:1000; #11229-1-AP; ProteinTech), p-MAP4 (1:1000, A51201; antibodies.com), CLIMP63 (CKAP4, 1:1,200, A302-257A; Fortis Life Science), AQP2 (1,1000, #3487; CST). HRP-conjugated secondary antibodies were applied (Immun-Star Goat Anti-Mouse, 1:20,000, 1705047 and Immun-Star Goat Anti-Rabbit, 1:20,000, 1705046, Biorad), and then activated with Clarity Max western ECL (Bio-Rad). Chemiluminescence was imaged with a ChemiDoc MP imager (Bio-Rad); signal intensity of each protein was measured with ImageLab software (Bio-Rad, v. 6.0) and normalized to levels of ⍺-tubulin or total protein (Ponceau S). At least 3-5 biological replicates were analyzed.

### Immunofluorescence microscopy

Cells were grown on coverslips and cultured in media for 24 hours. Cells subjected to osmotic stimulus were incubated in the specified osmotic buffer for 5-10 min and immediately fixed. Cells were fixed in 2% PFA for 20 min at room temperature, washed three times in phosphate-buffered saline (PBS), and blocked for 45 min in 2% BSA blocking solution with 0.2% triton X. The cells were incubated with antibodies against PC2 YCE2 (1:100, sc-47734, Santa Cruz Biotechnology), MAP4 (1:100; #SC-390286; Santa Cruz), CLIMP63 (CKAP4, 1:2,000, A302-257A; Fortis Life Science), AQP2 (1,1000, #3487; CST), Arl13b (1:100, #17711-1-AP; ProteinTech) overnight at 4°C, followed by the appropriate secondary antibody (Alexa Fluor 488 donkey anti-mouse IgG (1:1000, A21202, Invitrogen), Alexa Fluor 546 donkey anti-rabbit IgG (1:800, A10040, Invitrogen), for 1 hr, then washed three times in PBS. Some slides were co-incubated with 647-phalloidin (1:1000, 20555, Cayman Chemicals) during the secondary incubation step. Coverslips were mounted with Prolong-Diamond mounting media with DAPI (Invitrogen). After curing, cells were imaged with a 43X oil (N.A. 1.2) or 63X oil (N.A. 1.4) objective on an 880 Zeiss laser-scanning microscope with Airyscan (Zeiss, Germany). Images were post-processed with Zen Black software (Zeiss, Germany) and FIJI (NIH) [25].

### Live-cell imaging of microtubules and plus-end microtubules

Cells were plated on glass coverslip 48 hours prior to imaging. Imaging of microtubules was performed by staining the microtubules with ViaFluor Live Cell Microtubule Stain 647 (Biotium) following the manufacturer’s protocol. Tracking of the plus-end side of the microtubules was performed by transiently transfecting the cells with EB3-tdTomato (Addgene #50708) 24 hours after plating following the protocol described in the previous section. Cells were imaged with a 63x oil objective in an 880 Zeiss laser-scanning microscope with Airyscan.

### Analysis of live-cell imaging

Calcium imaging movies were analyzed using FIJI and the Time Series Analyzer V3 plugin. Quantification of the velocity, duration, and length of EB3-tdTomato comets was performed using the MtrackJ plugin in FIJI [26].

### Immunoprecipitation and Mass spectrometry

Protein (100 μg) isolated from the kidney (cortical and medullary) and left ventricle of wild-type C57 Bl6 mice was incubated with PC2 antibody (10 ug, Alomone laboratories) overnight complexed to magnetic protein G beads (Biorad). Following 3 rounds of washing, the bound protein was eluted and run, alongside flow-through on 4-12% polyacrylamide gels. Silver stain (Biorad) was used to identify bands for excision. Gel bands were excised and desiccated under hivac. Gel fragments (corresponding to 20, 30, 40, 70, 110, 200 and 250 kDa) were sent for LC-MS/MS analysis using an LTQ Orbitrap Elite at Rosalind Franklin University (Facility run by Dr. Charlie Yang). In-gel tryptic digestion was followed by the identification of proteins with the UniProt mouse database and PEAKS 8.5 software. Data were filtered based on −10^lgP^, FDR, unique peptide, and de novo ALC%. As an internal control, six bovine standard proteins were identified with high scores under the same conditions, to ensure that LC-MS/MS was running well.

### Statistical analysis

Data was plotted using GraphPad PRISM 9. Data were tested for normality using a normality and distribution test. For parametrically distributed data comparing two independent groups student’s t-test was used. For nonparametric data, Mann-Whitney analyses were conducted. Comparison of multiple groups with parametric distribution were analyzed through One or Two-Way ANOVA followed Sidak’s multiple comparison test. Comparison of multiple groups with non-parametric distribution were analyzed through One or Two-Way ANOVA followed by Kruskal-Wallis test. Conditions were considered statistically significant when *p* values were < 0.05. For live imaging experiments, quantifications were conducted using 10-20 cells per experimental condition from 3-5 biological replicates. Error bars indicate Standard Error of the Mean (SEM).

## Results

### Polycystin 2 tubule specific knock out mice have decreased urine osmolarity and impaired calcium response to hyperosmotic stimuli

ADPKD patients have decreased ability to concentrate urine, however, it is unclear if this defect is due to renal cysts or a signaling pathway impacted by the loss of functional polycystin proteins [6]. At a stage when cyst burden was not high [22], we analyzed whether urine osmolarity (i.e. urine concentration) was impaired in adult mice with induced PC2 kidney tubule-specific knock out (PC2 KO) versus the CTL (Fig. 1A). Two months after cre induction with doxycycline, the urine osmolarity of PC2 KO mice with *ad lib* water access was significantly reduced compared to CTL mice (Fig. 1B). Analysis of drinking, food intake, bodily free fluid, and energy expenditure did not reveal any differences between the CTL and PC2 KO mice (Fig. S1A-D) leading to the conclusion that decreased urine osmolarity was not due to changes in water and food consumption or the presence of renal cysts.

**Figure 1.**
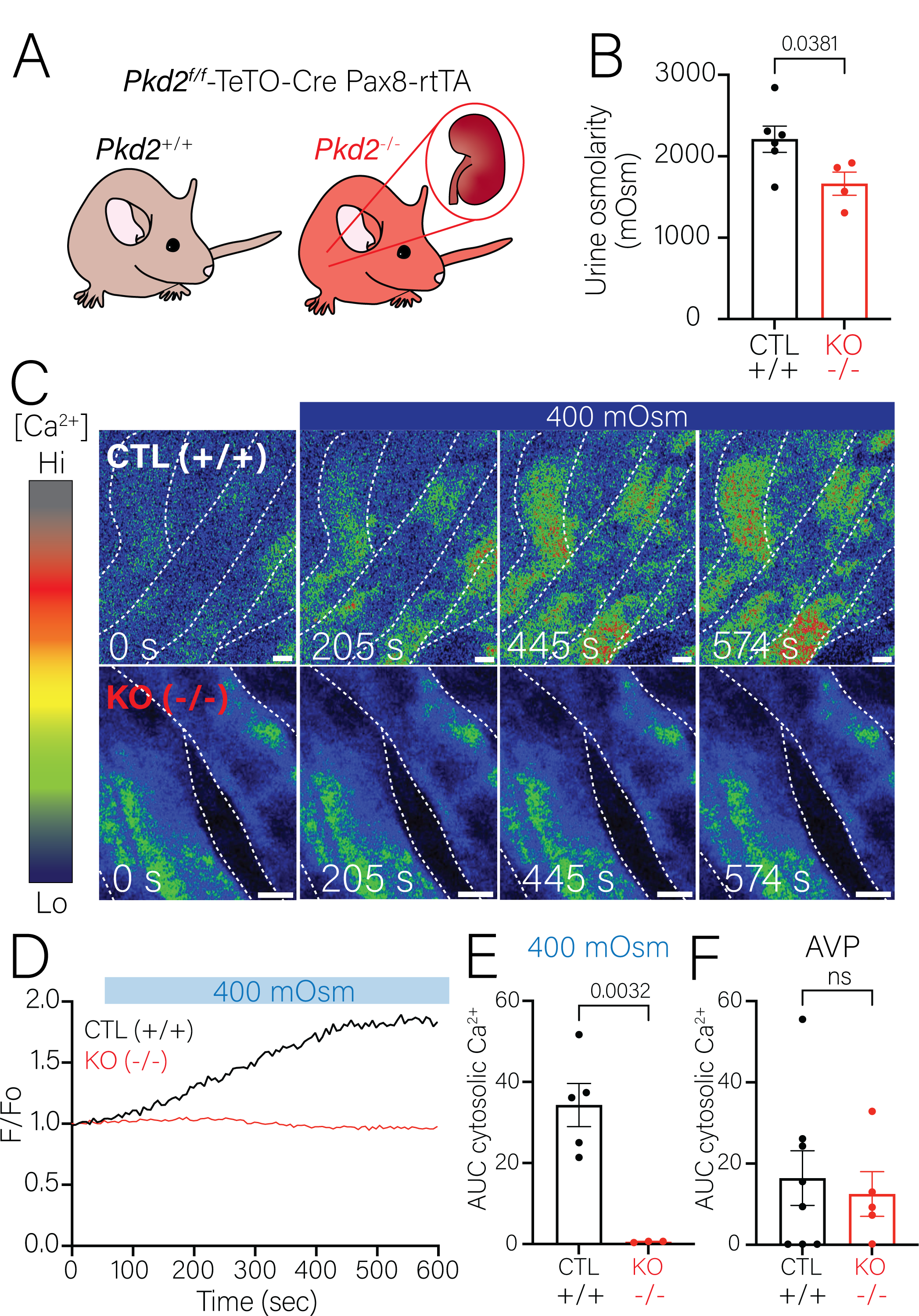
Tubule specific PC2 deletion leads to decreased urine concentration and decreased calcium signaling upon osmotic challenge in renal tubules. **A.** Model of tubule specific deletion of PC2 by crossing Pax8-rtTA Tet-O-Cre (CTL) mice with *Pkd2*^f/f^ mice. **B.** Deletion of tubule specific PC2 (Pax8-rtTA TetO-Cre *Pkd2*^f/f^ mice) resulted in decreased urine osmolarity in comparison to the CTL mice. Each dot represents an individual mouse. Bars represent mean±SEM. Data were analyzed to determine normality followed by student’s t-test to determine statistical analysis. p-values included in the figure. **C.** Representative time lapse images of collecting duct tubules from kidney slices incubated with Fluo-4 of CTL mice (top) and tubule specific PC2 KO mice (bottom). Extracellular osmolarity was increased from 250 mOsm to 400 mOsm. Scale bars represent 20 μm. **D.** Representative trace of cytosolic calcium changes in collecting duct tubules represented in panel *C.* Cytosolic calcium increased in CTL collecting duct tubules (*black line*) but not in PC2 KO collecting duct tubules (*red line*). **E.** Quantification of area under the curve was significantly decreased in PC2 KO collecting duct tubules in comparison to the CTL tubules. Bars represent mean±SEM. Data were analyzed to determine normality followed by student’s t-test. p-values are listed in the figure. **F.** Quantification of area under the curve was similar after stimulation with AVP in both CTL and PC2 KO collecting duct tubules. Bars represent mean±SEM. Data were analyzed to determine normality followed by student’s t-test.

Extracellular osmolarity is known to induce increase of cytosolic calcium [27, 28] and deletion of PC2 impairs calcium signaling in renal cells [29]. However, it has not been shown whether shifts in extracellular osmolarity stimulate PC2-mediated calcium release. To test if hyperosmotic shifts cause calcium release via a PC2 pathway, we tracked changes to cytosolic calcium of kidney sections in Salsa mice or incubated with the calcium sensor Fluo-4AM. An extracellular osmolarity shift (250 mOsm to 400 mOsm) in renal tubules from CTL mice stimulated an increase of cytosolic calcium (Fig. 1C-D, Video 1). In contrast, tubules from the PC2 KO mice had no increase in intracellular calcium after hyperosmotic stimuli (Fig. 1C-D). Note that the tubules from the PC2 KO mice were not cystic, and only showed mild dilation. Therefore, we consider these mice “pre-cystic” [22]. Quantification of the area under the curve (AUC) was significantly decreased in the tubules of the PC2 KO mice in comparison to the CTL (Fig. 1E). We identified medullary collecting duct tubules based on anatomical landmarks and ensured that these tubules responded to exogenously applied vasopressin in both CTL and PC2 KO mice (Fig. 1F).

### Deletion of Polycystin 2 abolished calcium signaling to osmotic changes

To dissect the role of PC2 in the osmosensing pathway in renal epithelial cells, we generated PC2 KO immortalized murine collecting duct cells (imCD3) using the CRISPR/Cas9 system. Validation of the PC2 deletion in the imCD3 cells was demonstrated with western blot (Fig. S2A-B) and immunofluorescence assay (Fig. S2C). Then both CTL and PC2 KO imCD3 cells were transfected with the genetic calcium reporter gCaMP6F to test whether PC2 was required for the cytosolic calcium signal after hyperosmotic challenge. We perfused cells with isosmotic solution (300 mOsm) for 3 minutes (at 3ml/min) to ensure there was no contribution of fluid flow or sheer stress. Extracellular osmolarity was then increased to 400 mOsm in both cell lines and the flow rate kept the same. In CTL cells, the hyperosmotic stimulus induced an increase of cytosolic calcium, which was absent in the PC2 KO cells (Fig. 2A, Video 2). Representative tracings of cytosolic calcium changes in both CTL and PC2 KO cells are shown in Fig. 2B and Video 2. Quantification of area under the curve and peak cytosolic calcium were significantly decreased in the PC2 KO cells (Fig. 2C-D).

**Figure 2.**
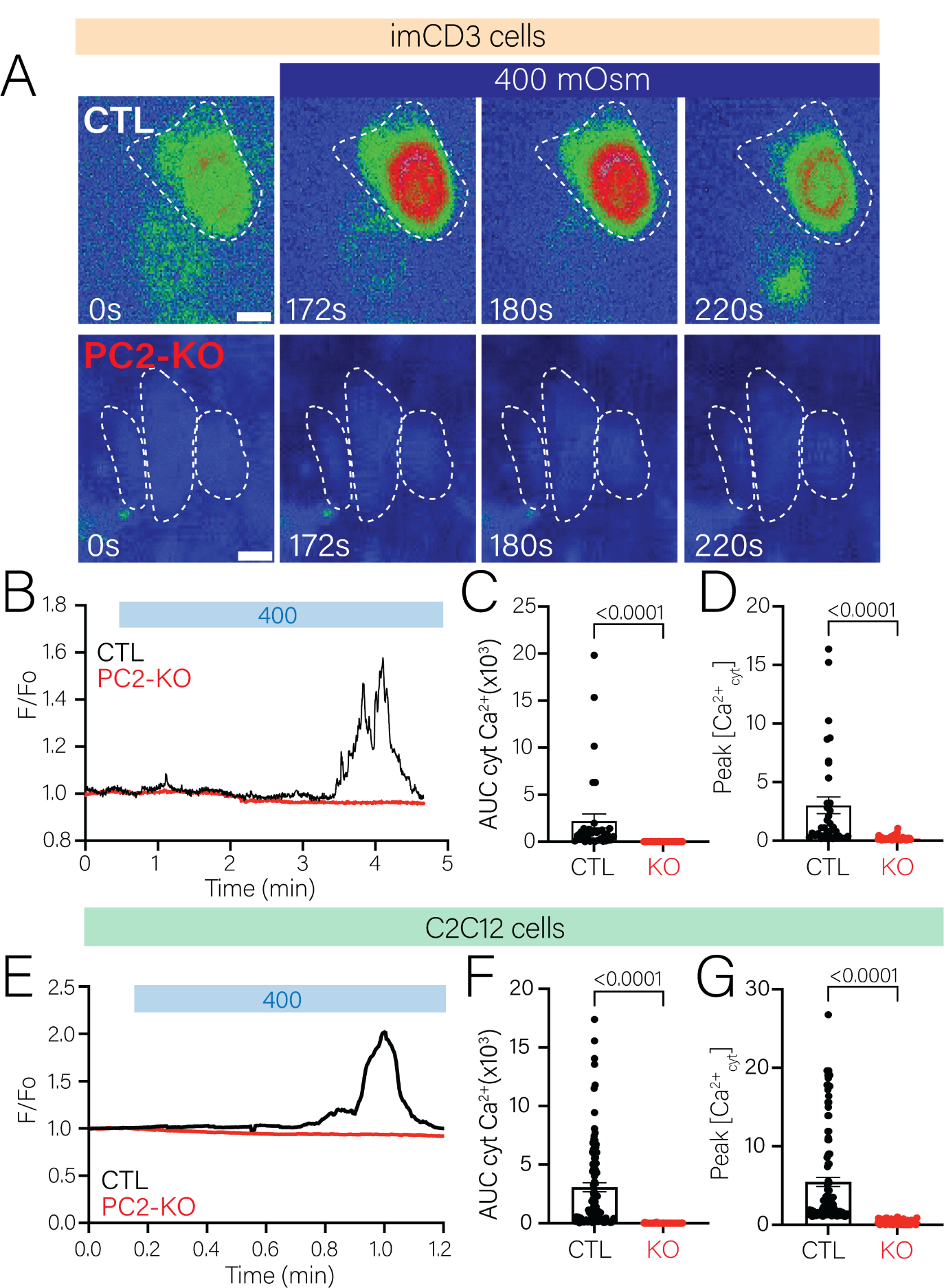
Deletion of PC2 in imCD3 and C2C12 cells abolished cytosolic calcium increase upon increase of extracellular osmolarity. **A.** Representative time lapse images of CTL and PC2 KO imCD3 cells transfected with cytosolic gCaMP6F demonstrating absense of calcium signal in PC2 KO cells when extracellular osmolarity was increased from 300 mOsm to 400 mOsm. Scale bars represent 10 μm. **B.** Representative analysis of cytosolic calcium changes in CTL (*black line*) and PC2 KO imCD3 cells (*red line*). Cytosolic calcium increase was abolished in PC2 KO cells. **C.** Area under the curve was significantly decreased in PC2 KO imCD3 cells. Bars represent mean±SEM. Data were first analyzed to determine normality then followed with a Mann-Whitney’s statistical test. p-values listed in the figure. **D.** Peak cytosolic calcium was significantly decreased in the PC2 KO imCD3 cells. Bars represent mean±SEM. Data were analyzed to determine normality followed by Mann-Whitney’s statistical test. p-values listed in the figure. **E.** Representative graph of cytosolic calcium changes in CTL (*black line*) and PC2 KO C2C12 cells (*red line*). Cytosolic calcium increase was absent in PC2 KO cells. **F.** Area under the curve was significantly decreased in C2C12 PC2 KO cells. Bars represent mean±SEM. Data were first analyzed to determine normality then followed with a Mann-Whitney’s statistical test. p-values listed in the figure. **G.** Peak cytosolic calcium was significantly decreased in the C2C12 PC2 KO cells. Bars represent mean±SEM. Data were first analyzed to determine normality then followed with a Mann-Whitney’s statistical test. p-values listed in the figure.

To demonstrate that the hyperosmotically induced cytosolic calcium increase was indeed mediated by PC2, and were reproducible in another cell line, we performed the same experiment in C2C12 myoblast cells, with PC2 knocked out [30]. We observed that like the renal tubules and the imCD3 cells, the C2C12 CTL cells had increased cytosolic calcium in response to extracellular osmotic changes (Fig. 2E, *black line*) whereas the response was absent in the C2C12 PC2 KO cells (Fig. 2E-G). To test that the cytosolic calcium increase was due to the osmotic pressure and not the salt gradient, we increased extracellular osmolarity with mannitol. This increase of extracellular osmolarity through mannitol still elicited a cytosolic calcium response in the C2C12 CTL cells which was absent in the PC2 KO cells (Fig. S3A-B). Thus, we found that the cytosolic calcium increase induced by hyperosmotic changes in the different tissues and cell lines analyzed was absent where PC2 was deleted (i.e. isolated kidney tubules, imCD3 and C2C12 myoblast cells). These results suggest that the calcium signaling induced by hyperosmotic changes is mediated by PC2. As the observed response was similar in the tubules and two cell lines analyzed, we continued to dissect the osmosensing mechanism in C2C12 cells to determine how ER localized PC2 senses extracellular changes.

### The osmotically induced calcium signal does not originate from the plasma membrane or cilia

PC2 has been reported to localize to the primary cilia, plasma membrane and the ER in different tissues [10]. To determine which localization the osmotically induced calcium signal originated from, we first conducted experiments in the absence of extracellular calcium. Even in the absence of extracellular calcium, an increase of extracellular osmolarity still elicited a calcium response in CTL cells which was absent in the C2C12 PC2 KO cells (Fig. S3C-D). These results suggest that the calcium response due to osmotic changes comes from intracellular calcium release and not calcium entry. We then dissected the kinetics of the calcium responses in different cellular compartments where PC2 is known to reside, namely the ER, the plasma membrane, and the primary cilia using a variety of genetically encoded calcium indicators (Fig. 3A). We co-expressed R-cepia (ER calcium indicator) and gcamp7s-CAAX (a plasma membrane calcium indicator) to simultaneously measure changes in the ER and plasma membrane. Hyperosmotic stimulus first induced a reduction in ER calcium (R-cepia) indicating calcium release from the ER (Fig. 3B-C, region “a”), subsequently followed by an increase in plasma membrane calcium (Fig. 3B-C, region ‘b”). This observation suggests that hyperosmotic changes induced ER calcium release which subsequently stimulated calcium influx. Consistent with the previous finding, in the presence of extracellular calcium, CTL cells elicited plasma membrane calcium sparks which were absent in the C2C12 PC2 KO cells (Fig. 3D). The lack of plasma membrane calcium sparks in the C2C12 PC2 KO cells is interpreted to be due to the inability to activate calcium activated calcium influx upon hyperosmotic changes.

**Figure 3.**
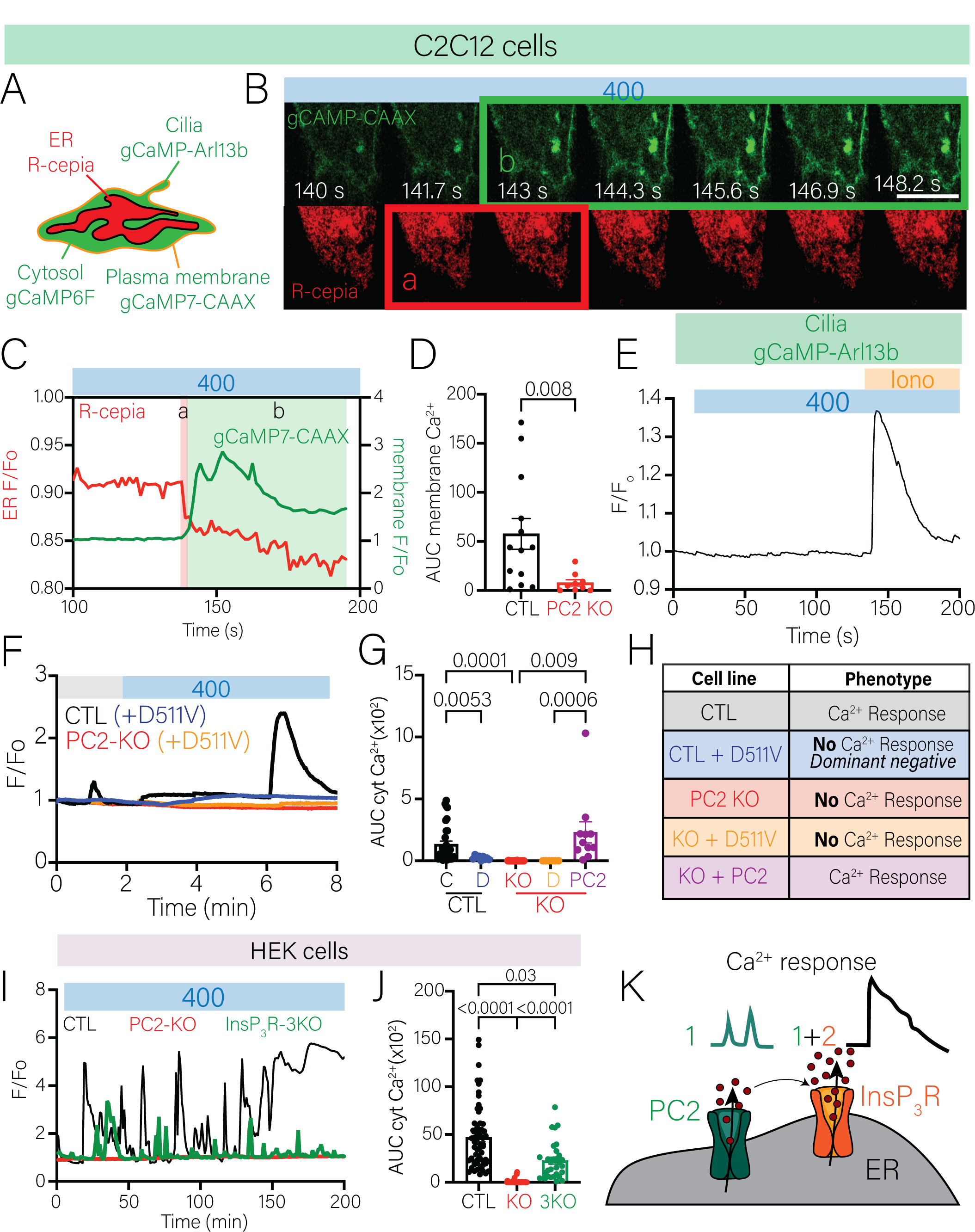
Calcium mediated by PC2 in the ER is essential for the hyperosmotic response and recruits InsP3R to sustain cytosolic calcium. **A.** Diagram highlighting the different genetic calcium indicators used for the cilia, plasma membrane, ER and cytosol. **B.** Representative images of C2C12 CTL cells expressing the plasma membrane calcium indicator (gCaMP7s-CAAX, green, top panels) and ER calcium indicator (R-cepia, red, bottom panels). Green and red box represents calcium changes graphed in panel *F*. Scale bars represent 10 μm. **C.** Representative trace of simultaneous calcium changes in R-cepia (red line) and plasma membrane calcium (green) in C2C12 CTL cells after hyperosmotic stimulus. Red box indicates ER calcium drop after hyperosmotic stimuli highlighted in panel *B*. Green box indicates plasma membrane calcium spark highlighted in panel *B*. **D.** Membrane calcium area under the curve was significantly decreased in PC2 KO cells. Bars represent mean±SEM. Data were analyzed to determine normality. Statistical analysis was determined by student’s t-test. p-values are listed in the figure. **E.** Ciliary calcium did not increase in CTL C2C12 cells after increase of extracellular osmolarity (400 mOsm) but did increase when Ionomycin (1μM) was added. **F.** Representative graph of cytosolic calcium changes in C2C12 CTL (*black line*), C2C12 CTL + D511V PC2 variant (*blue line*), C2C12 PC2 KO (*red line*), C2C12 PC2 KO + D511V PC2 variant (*orange line*) cells. **G.** The D511V variant acts as a dominant negative in C2C12 CTL cells and expression of full-length PC2 restored increase of cytosolic calcium in the PC2 KO cells. Bars represent mean±SEM. Data were analyzed to determine normality. Statistical analysis was determined by a Two-way ANOVA test followed by Sidak’s. p-values listed in figure. **H.** Table summarizing the cell lines analyzed and phenotype observed graphed in panels *F-G*. **I.** Representative trace of cytosolic calcium changes in HEK CTL (*black line*), HEK PC2 KO (*red line*) and 3KO-HEK (*green line*) cells. Cytosolic calcium increase was absent in PC2 KO cells. **J.** Area under the curve was significantly decreased in both HEK PC2 KO and 3KO-HEK cells. Bars represent mean±SEM. Data were analyzed to determine normality. Statistical analysis was determined by One-way ANOVA followed by Kruskal-Wallis analysis (Gaussian distribution was not assumed). p-values listed in figure. **K.** Hyperosmotic stimuli requires an initial calcium release mediated by PC2 which is further sustained by recruiting InsP_3_R localized in the ER.

A substantial amount of previous work has demonstrated that PC2 localizes to primary cilia, where it can participate in calcium influx [14–16, 18]. We therefore asked whether osmotic changes were inducing a ciliary response amplified by the cytosolic response. Using an Arl13B antibody we quantified the percentage of cells with cilia and found that there was no significant difference between C2C12 CTL and PC2 KO cells (Fig. S4A-B). We then used a ciliary calcium indicator, gCaMP6f-Arl13B, and found that osmotic stimuli did not stimulate ciliary calcium response (Fig. 3E and Fig. S4C). We ensured that the ciliary sensor is functional by applying ionomycin after the hyperosmotic stimuli and observed robust ciliary calcium increases (Fig. 3E). Taken together, we demonstrated that hyperosmotic stimuli induced ER-localized PC2 calcium release which further activates calcium influx from the plasma membrane to maintain the cytosolic calcium response. These experimental results strongly suggest that neither ciliary PC2 nor plasma membrane localized PC2 contributes to the initial osmotically driven calcium response.

### Expression of the PC2 channel dead mutant acts as dominant negative in control cells

To validate that the initial calcium response is mediated by PC2, we over-expressed full length PC2 and a pathogenic variant (D511V) that does not allow ion flux through PC2. Expression of the PC2-D511V variant in the CTL cells abolished the calcium response, demonstrating that the mutation acted as a dominant negative (Fig. 3F, *blue line*). Expression of PC2-D511V in the PC2-KO cells did not restore the calcium response upon hyperosmotic stimuli (Fig. 3F, *orange line*). Importantly, over-expression of the full length PC2 restored the calcium response (Fig. 3G). Quantification of the AUC is shown in Figure 3G for each of the conditions (summary of response shown in Fig. 3H). These data suggest that the osmotically induced calcium response requires not only the expression of PC2, but also a functional ion conducting PC2 to activate calcium release.

### Polycystin 2 recruits InsP_3_R to sustain increase of cytosolic calcium due to osmotic stimuli

Although the previous experiments suggested that the calcium signal originated from ER-localized PC2, it seemed unlikely that PC2 was acting by itself to sustain the calcium release. Previous work has demonstrated that PC2 can regulate the activity of other ER-localized calcium channels like the InsP_3_R [11–13].To determine if the lack of observed calcium response was due to the absence of an interaction between PC2 and InsP_3_R, we used a cell line, with the three isoforms of the InsP_3_R knocked out (hereby called 3KO-HEK generated from Human Embryonic Kidney (HEK) cells) [23]. We compared the HEK cells with PC2 KO we generated via CRISPR/Cas9 and validated through western blot and quantitative real-time PCR (RT-PCR) (Fig. S5A-C). After increasing extracellular osmolarity, we observed a reduced cytosolic calcium increase in the 3KO-HEK cells (Fig. 3I-J). Quantification of the AUC in the 3KO-HEK cells showed there was ∼50% less compared with the HEK CTL cells (Fig. 3J). Moreover, the osmotically induced response in the HEK CTL cells was composed of two different types of calcium peaks: a sharp and fast response (*green line* in Fig. 3K) and a sustained cytosolic calcium response (*black line* in Fig. 3K). However, only the sharp and fast response (*green line*) was observed in the 3KO-HEK cells. These data suggest that the osmotic stimuli induce an initial calcium release mediated by ER-localized PC2 then further recruited calcium channels, like the InsP_3_R, to sustain the cytosolic calcium response (Fig. 3K).

### Microtubules aid in mediation of the osmosensing response

The data presented indicates that ER-localized PC2 initiates the calcium increase in response to changes in extracellular osmolarity. These data raise the question, how is an external osmotic stimulus being transmitted to the ER? Osmolarity induces volumetric changes that are mediated by rapid changes to the cytoskeleton [31]. Microtubules directly link the cytoskeleton to the ER [32]. To determine if the microtubules mediate the osmotic response, we tested the effects of the microtubule inhibitors paclitaxel (Taxol) and nocodazole on the osmotic response (Fig. 4A). Taxol stabilizes microtubules, preventing depolymerization [33], whereas nocodazole induces depolymerization of the microtubules, causing destabilization and collapse of the microtubules [34]. We incubated the cells with the microtubule inhibitors in isosmotic solution (300 mOsm) and increased extracellular osmolarity with the inhibitors. Application of the isosmotic solution with nocodazole (5μM) induced an initial calcium increase (Fig. 4B, turquoise line), which we interpret as the ER-localized PC2 calcium release upon depolymerization of the microtubules. However, upon increase of extracellular osmolarity, there was no rise in the cytosolic calcium in the presence of either nocodazole or Taxol (Fig. 4B,D). The absence of the cytosolic calcium response was due to prevention of the microtubules to relay the extracellular changes; this was confirmed as the addition of thapsigargin (inhibitor of the sarcoendoplasmic reticulum calcium ATPase (SERCA) which allows leakage of the ER into the cytosol) in the presence of hyperosmotic stimuli induced calcium release (Fig. 4C). These data suggest that the ability to release intracellular calcium after hyperosmotic stimuli is coupled to the integrity of the microtubules.

**Figure 4.**
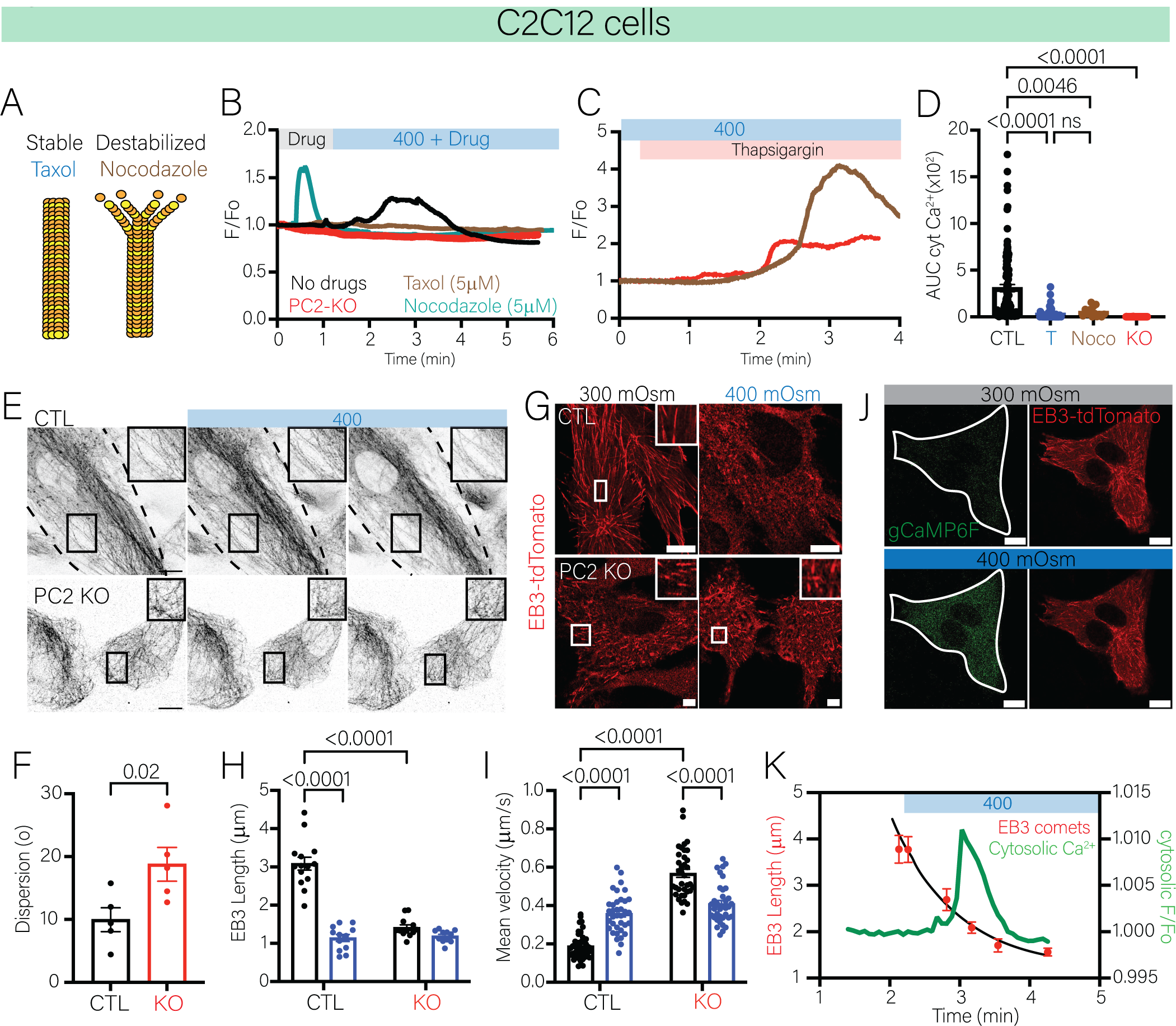
Cytosolic calcium response induced by osmotic stimuli is mediated by interaction between the microtubules and ER. **A.** Mode of action of microtubule inhibitors used in panel *B*. Paclitaxel (Taxol) promotes stabilization of microtubules in comparison to nocodazole which promotes destabilization/depolymerizing of microtubules. **B.** Representative trace of cytosolic calcium changes in C2C12 CTL cells with no drugs (*black line*), C2C12 CTL + nocodazole (*brown line*) and C2C12 PC2 KO (*red line*) cells. **C.** Representative trace of cytosolic calcium increase after the addition of thapsigargin (1μM) in C2C12 PC2 KO cells and CTL cells with Paclitaxel. **D.** Area under the curve was significantly decreased in C2C12 CTL cells incubated with both Paclitaxel and nocodazole which resembles the response in the C2C12 PC2 KO cells. Bars represent mean±SEM. Data were analyzed to determine normality. Statistical analysis was determined by One-way ANOVA followed by Kruskal-Wallis (Gaussian distribution was not assumed). p-values listed in the figure. **E.** Representative images of C2C12 CTL cells (top panels) and PC2 KO cells (bottom) labeling microtubules at 300 mOsm and after hyperosmotic stimuli (400 mOsm). **F.** Quantification of the degree of dispersion of the microtubules at 300 mOsm of C2C12 CTL and PC2 KO cells. **G.** Representative images of EB3-tdTomato expression in C2C12 CTL cells at 300 mOsm and 400 mOsm (top panels) and C2C12 PC2 KO cells under the same conditions (bottom panels). *Insets*: smaller EB3 comets in the C2C12 PC2 KO cells under 300 mOsm (*black bars*) and 400 mOsm (*blue bars*). Scale bars represent 10 μm. **H.** EB3 comet length decreased in C2C12 CTL after addition of 400 mOsm (*blue vs black bars*). Comets were significantly reduced in basal osmolarity (300mOsm; *black bars*) and remained unchanged at 400mOsm (*blue bars*) in PC2 KO cells. Bars represent mean±SEM. Data were analyzed to determine normality. Statistical analysis was determined by Two-way ANOVA followed by Sidak’s test. p-values listed in the figure. **I.** Mean velocity of EB3 comets was increased upon increase of extracellular osmolarity (*blue bar vs black bar*). EB3 comets were significantly faster in PC2 KO cells at basal osmolarity (300mOsm; *black bars*) and decreased after hyperosmotic stimuli (400 mOsm; *blue bars*), although still faster than the CTL cells. Bars represent mean±SEM. Data were analyzed to determine normality. Statistical analysis was determined by Two-way ANOVA followed by Sidak’s test. p-values listed in the figure. **J.** Representative images of C2C12 CTL cells expressing a cytosolic calcium indicator and EB3-tdTomato at basal osmolarity (*top panels*) and at 400 mOsm (bottom panels). Scale bars represent 10 μm. **K.** Shortening of EB3 comets in CTL cells (red dots) occurs within 0.91 min after hyperosmotic stimuli which induced a cytosolic calcium (green trace) increase 20 seconds after. Errors bars represent SEM.

To assess the integrity of microtubules and determine if hyperosmolarity caused depolymerization of the microtubules, we stained the microtubules with ViaFluor 647. At baseline conditions (300 mOsm), we observed that the structure of the microtubules was different in C2C12 CTL versus PC2 KO cells (Fig. 4E). Microtubules in the CTL cells were linear and stretched across the whole cell whereas in the PC2 KO cells, microtubules were dispersed and disorganized (Fig. 4E, first panel of CTL and PC2 KO cells). The degree of dispersion of the microtubules throughout the cells was significantly increased in the PC2 KO cells (Fig. 4F). A higher degree of dispersion is interpreted as less directional and more dispersed distribution from the microtubules throughout the cell. Increase in extracellular osmolarity caused retraction of the microtubules in the CTL cells (Fig. 4E, *inset top panels*). However, in the PC2 KO cells microtubules remained stable after hyperosmotic change (Fig. 4E, *inset bottom panel*).

We then examined microtubule growth dynamics by expressing EB3-tdTomato which binds to the plus-end of microtubules (Fig. 4G, Video 3). In CTL cells, the length of EB3 comets in isosmotic conditions was approximately 3 µm (Fig. 4H, *black bars in CTL cells*). In CTL cells under hyperosmotic stimulus, the length of the EB3 comets was decreased to ∼ 1.5 µm (Fig. 4H, *blue bars in CTL cells*). In PC2 KO cells, EB3 length under isosmotic conditions was significantly shorter (∼1.5 µm) and remained the same length under hyperosmotic conditions (Fig 4H). Additionally, the velocity of the EB3 comets under hyperosmotic conditions in CTL cells was significantly increased, indicating microtubule growth (Fig. 4I). In contrast, the mean velocity of EB3 comets was significantly increased at 300 mOsm but decreased upon hyperosmotic conditions in PC2 KO cells (Fig. 4I). EB3 comet dynamics for C2C12 CTL and PC2 KO cells before and after hyperosmotic changes are described in Table 1.

**Table 1.** EB3 kinetics in C2C12 CTL and PC2 KO cells.

| Parameters | Cell lines | 300 mOsm | 400 mOsm |
| --- | --- | --- | --- |
| Average length ($\mu\text{m}$) | CTL | $3.08 \pm 0.65$ | $1.13 \pm 0.32$ |
| | PC2 KO | $1.40 \pm 0.24$ | $1.18 \pm 0.16$ |
| Average velocity ($\mu\text{m/s}$) | CTL | $0.18 \pm 0.06$ | $0.35 \pm 0.10$ |
| | PC2 KO | $0.56 \pm 0.06$ | $0.40 \pm 0.06$ |
| Duration of comet (s) | CTL | $32.26 \pm 9.88$ | $9.49 \pm 4.52$ |
| | PC2 KO | $9.28 \pm 2.52$ | $10.20 \pm 3.42$ |

To dissect the coupling of the microtubule rearrangement to the calcium signal induced by hyperosmolarity, we co-expressed gCaMP6F and EB3-tdTomato, to simultaneously measure cytosolic calcium increase and microtubule dynamics (Fig. 4J). Following the hyperosmotic change, the length of the EB3 comets quickly decreased to ∼1.5 µm (Fig. 4F). The decreased length of the EB3 comets was taken to indicate microtubule collapse, which was then followed by an increase of cytosolic calcium (Fig. 4F, *red dots vs green line*). These data strongly suggest that hyperosmolarity first induces microtubule depolymerization, which then leads to intracellular calcium release from ER localized PC2. The absence of calcium signals from PC2 KO cells is likely due to the combined effect of impaired microtubules organization and dynamics leading an inability to relay external stimuli to intracellular signals like calcium release.

### PC2 interacts with microtubule components

To identify how PC2 interacts with microtubules in the osmosensitive pathway, we performed immunoprecipitation assays and mass spectrometry with PC2 from kidney tissue. The identified list of proteins included new potential interactors such as Microtubule Associated Protein (MAP) 4 (Supp. Table 1). We focused on MAP4 as it is known to bind to microtubules to promote stabilization [35]. We validated the reciprocal interaction between PC2 and MAP4 by immunoprecipitation in C2C12 myoblast cells (Fig. S6A). Staining of PC2 and MAP4 in C2C12 CTL cells showed partial colocalization of these proteins (Fig. S6B). We also stained CTL cells and PC2 KO cells to determine the distribution of MAP4 in the cells. We observed that in PC2 KO cells, distribution of MAP4 appears evenly distributed throughout the cell in comparison to the CTL cells (Fig. S6C). Localization of MAP4 in the CTL cells closely resembled the distribution of the ER, labeled with CLIMP63 (Fig. S6D).

### Deletion of PC2 affects ER morphology

Association of the microtubules with organelles [36], including the ER, leads to coordinated movement of MTs and ER [36]. The association formed by the MT-ER can also dictate and regulate ER morphology [32]. As such, we explored whether the disruption of the PC2-MAP4 interaction affected ER morphology. In comparison to the CTL cells, the PC2 KO cells had a higher ratio of cisternae to tubules (Fig. S7A-B). Higher cisternae ratio indicates a less motile and less dynamic ER. We examined CLIMP63 expression to visualize the ER in fixed cells (Fig. S7C). In CTL cells, the ER was spread throughout the cells, whereas the ER was confined to the nuclear area in PC2 KO cells. Additionally, CLIMP63 particle size was significantly lower in PC2 KO cells, suggesting that the ER covered less surface area than in the CTL cells. (Fig. S7D). Re-expression of full-length PC2 in the C2C12 PC2 KO cells restored the tubular conformation in the ER (Fig. S7E).

Deletion of PC2 collapsed and confines the ER to the nuclear area, therefore we quantified the area between the plasma membrane and the ER. We transfected ER-mCherry along with a gCaMP7s-CAAX membrane marker and observed that in PC2 KO, the space between the ER membrane and plasma membrane was significantly increased (Fig. S7F-G). Taken together, these data demonstrate that disruption of the tethering of the ER and the microtubules is mediated by PC2 and MAP4, and that this interaction not only regulates ER morphology but as shown in Fig. 3, microtubule dynamics. Moreover, the increased spacing between the ER and plasma membrane in PC2 KO cells means that under external stimulus PC2 contributes to more dynamic intracellular movement and adaptation by altering ER morphology.

### PC2 affects expression and phosphorylation of MAP4

Next, we aimed to ask how do PC2 and MAP4 mechanistically interact upon hyperosmotic stimuli. It has been shown that binding of MAP4 to microtubules promotes stabilization, and upon phosphorylation, MAP4 disassociates from the microtubules, which destabilizes and promotes depolymerization of the microtubules [37]. Consistent with the literature, we found that phosphorylation of MAP4 to total protein was significantly increased upon hyperosmotic change in CTL cells (Fig. 5A-B). In PC2 KO cells, phosphorylated MAP4, was increased in isosmotic conditions and remained unchanged after hyperosmotic change (Fig. 5A-B). After hyperosmotic stimuli, we observed that total MAP4 remained unchanged in CTL cells after 1 hour (Fig. 5C-D). In contrast, deletion of PC2 significantly decreased expression of total MAP4, to ∼50% less and remained unchanged upon hyperosmotic stimulus (Fig. 5C-D). To demonstrate that these changes were specific to the MAP4 pathway, we measured the expression of another protein which binds to microtubules and localizes to the ER, CLIMP63. Expression of CLIMP63 remained unchanged upon increase of extracellular osmolarity or deletion of PC2 (Fig. 5F).

**Figure 5.**
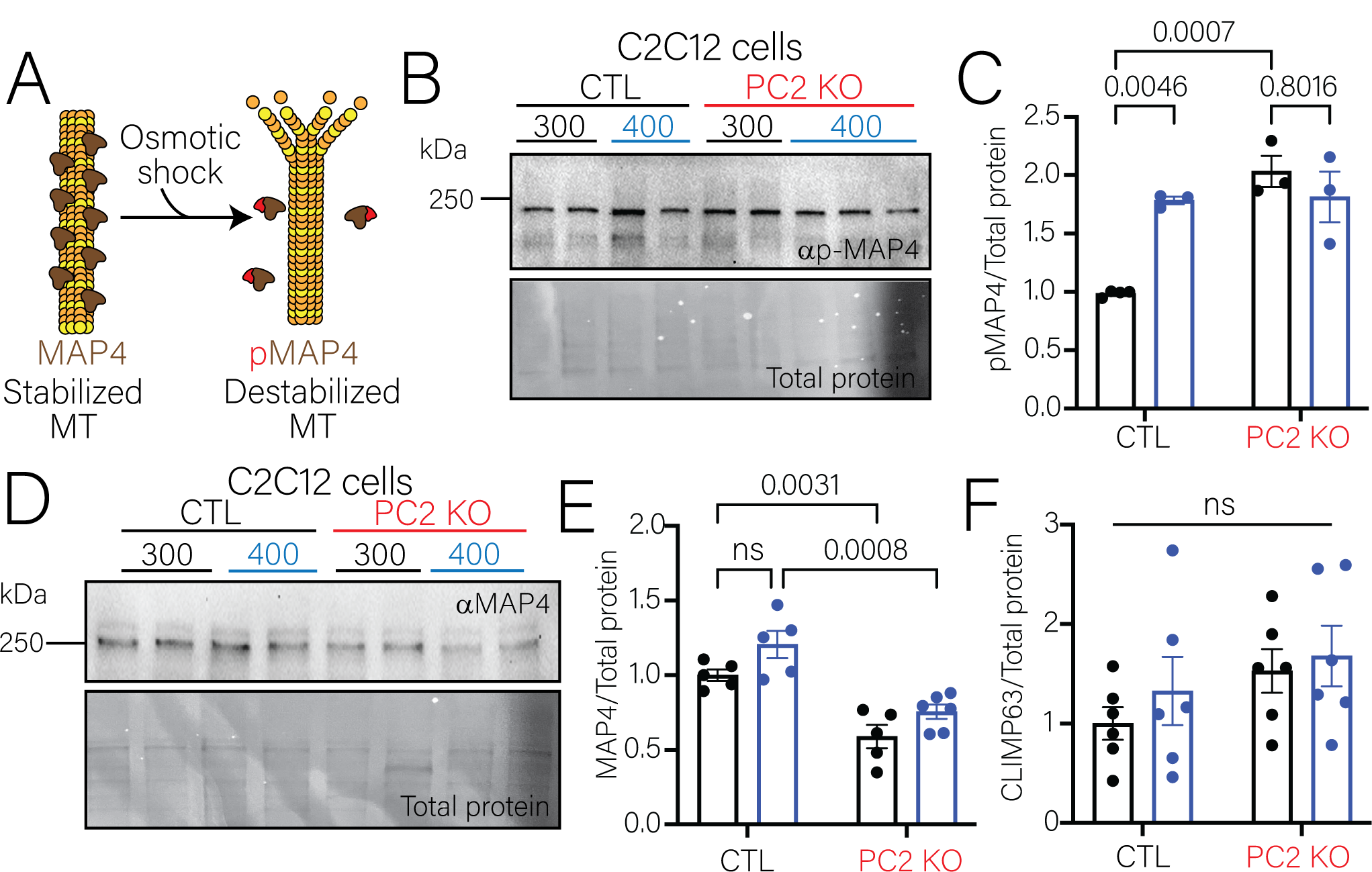
PC2 regulates MAP4 expression. **A.** MAP4 binds and stabilizes microtubules. Disassociation of MAP4 from the microtubules due to phosphorylation of MAP4 destabilizes microtubules. **B.** Expression of p-MAP4 in C2C12 CTL and PC2 KO cells at 300 mOsm and 400 mOsm. Total protein was used as loading control. **C.** p-MAP4 increased in C2C12 CTL cells after increasing extracellular osmolarity. Basal p-MAP4 levels were higher in PC2 KO cells compared to CTL but did not change with osmolarity. Bars represent mean±SEM. Data were analyzed to determine normality. Statistical analysis was determined by Two-way ANOVA followed by Sidak’s test. p-values listed in the figure. **D.** Western blot of MAP4 expression in C2C12 CTL and PC2 KO cells at 300 mOsm and 400 mOsm. Total protein was used as loading control. **E.** MAP4 expression was significantly decreased in PC2 KO cells. Bars represent mean±SEM. Data were analyzed to determine normality. Statistical analysis was determined by Two-way ANOVA followed by Sidak’s test. p-values listed in the figure. **F.** Expression of CLIMP63 remained unchanged in C2C12 CTL and PC2 KO cells. Bars represent mean±SEM. Data were analyzed to determine normality. Statistical analysis was determined by Two-way ANOVA followed by Sidak’s test.

To determine if the decreased expression of MAP4 was dependent on the calcium activity of PC2 or the interaction with PC2, we overexpressed WT PC2 and the calcium dead channel variant (D511V) in the PC2 KO cells. Under hyperosmotic stimulation, increase expression of MAP4 was restored in the presence of either the full-length PC2 or the D511 variant (Fig. S8A). These results suggest that upon increase of extracellular osmolarity MAP4 is phosphorylated and disassociates from the microtubules, leading to unstable microtubules. Depolymerization of the microtubules then “tugs” on PC2 allowing for PC2 mediated calcium release. Deletion of PC2 disrupts the connection between the microtubules and the ER through its association with MAP4, which further decreases the expression of MAP4.

### Deletion of MAP4 phenocopies PC2 KO cells

We then confirmed that an interaction between PC2 and MAP4 is critical in allowing the cell to mediate osmotic responses. Using CRISPR/Cas9, we transfected C2C12 cells with MAP4 specific guide RNA to generate a polyclonal knockout of MAP4. We validated that MAP4 expression was decreased 75% through western blot (Fig. 6A-B) and by immunofluorescence assay (Fig. 6C). Expression of PC2 in the MAP4-KO was significantly increased (Fig. S9A-B) but the localization of PC2 remained unchanged in the MAP4-KO cells (Fig. S9C). We examined CLIMP63 expression to determine if the MAP4-KO affected the distribution of the ER (Fig. S9C). In CTL cells, CLIMP63 was evenly distributed throughout the cell and co-localized with MAP4 (Fig. 6C). However, in the MAP4-KO cells we observed that the ER remained closer to the nuclear area, like the phenotype observed in the PC2 KO cells (Fig. 6C; Fig. S7C). Then, using the gCaMP6F, we monitored changes to the cytosolic calcium upon increase of extracellular osmolarity and found that deletion of MAP4 abolished the cytosolic calcium increase (Fig. 6D). AUC was significantly reduced in the MAP4-KO cells compared to CTL cells (Fig. 6E). Addition of thapsigargin increased cytosolic calcium in the MAP4-KO, indicating the cells were able to respond to calcium agonists (Fig. 6F). Earlier data suggested the deletion of PC2 affected the microtubules dynamics; we examined if MAP4-KO has a similar effect by measuring the length of EB3 comets at baseline and after hyperosmotic stimuli (Fig. 6G). In comparison to the CTL cells, the length of EB3 comets was significantly shorter in the MAP4-KO (Fig. 6H). Taken together, deletion of MAP4 phenocopied the PC2 KO cells in the osmotically induced response. These results further validate that the association of MAP4 and PC2 are responsible for the osmosensing pathway.

**Figure 6.**
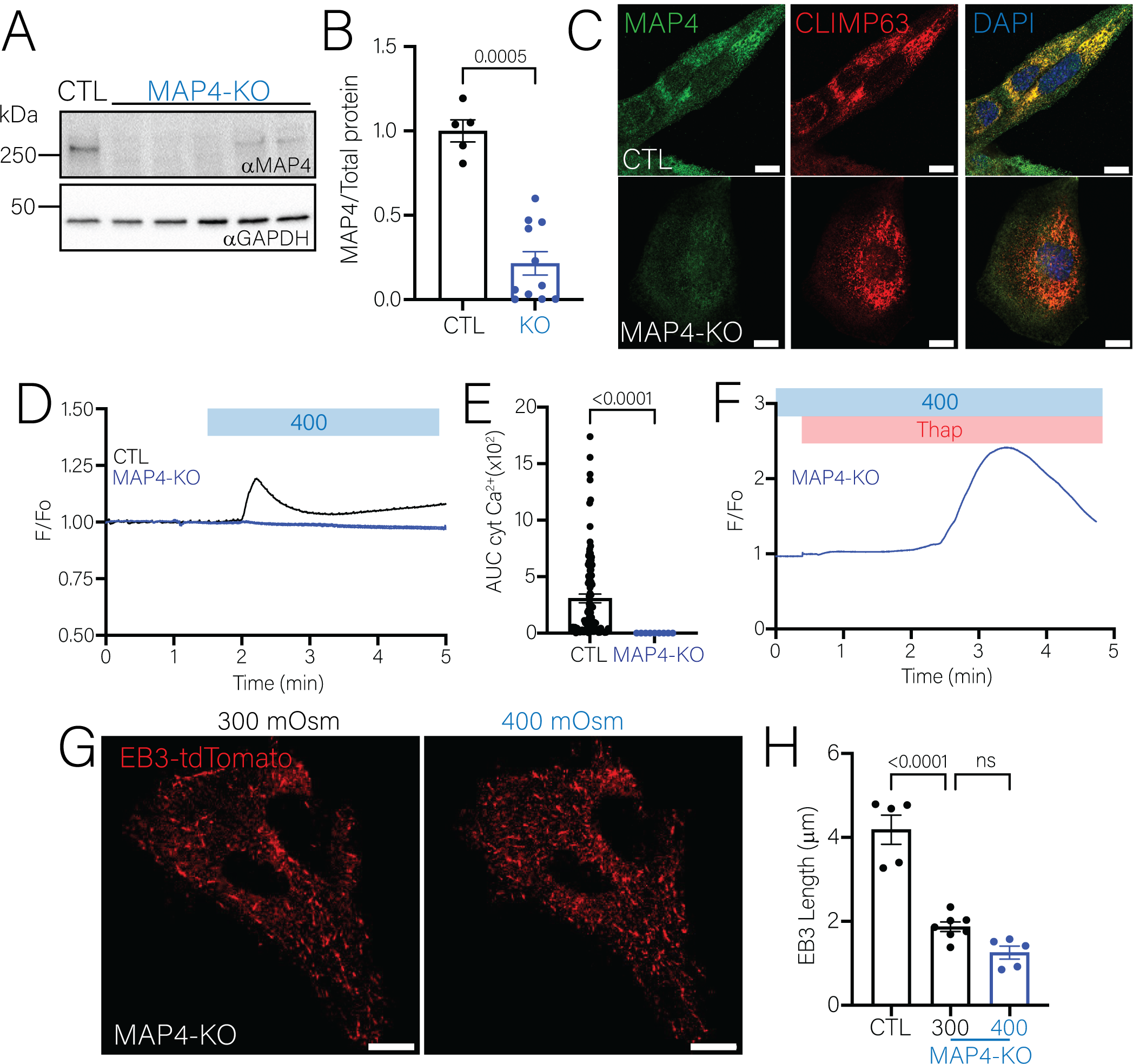
Knock out of MAP4 phenocopies osmosensitive response of PC2 KO cells. **A.** Expression of MAP4 in C2C12 CTL and MAP4-KO cells. GAPDH was used as loading control. **B.** MAP4 expression was decreased 75% in the knock down cell line. **C.** Immunofluorescence staining of MAP4 and CLIMP63 in C2C12 CTL cells (top panels) and MAP4-KO cells (bottom panels). **D.** Representative trace of cytosolic calcium changes in C2C12 MAP4-KO cells (*blue line*). Scale bars represent 10 μm. **E.** Area under the curve was significantly decreased in the MAP4-KO after increasing extracellular osmolarity. Bars represent mean±SEM. Data were analyzed to determine normality. Statistical analysis was determined by student’s t-test. p-values listed in figure. **F.** Representative trace of cytosolic calcium increase using thapsigargin after increasing extracellular osmolarity in MAP4-KO cells. **G.** Representative images of EB3-tdTomato expression in C2C12 MAP4-KO cells at 300 mOsm (left panel) and 400 mOsm (right panels). Scale bars represent 10 μm. **H.** EB3 comet length was significantly decreased in C2C12 MAP4-KO at basal osmolarity (300 mOsm) and remained further unchanged after osmotic increase. Bars represent mean±SEM. Data were analyzed to determine normality. Statistical analysis was determined by One-way ANOVA. p-values listed in figure.

### Biological implications of the PC2-MAP4 osmosensing pathway

Earlier in this study, we showed that the urine osmolarity in the kidney tubule specific PC2 KO mice were significantly decreased compared to the CTL mice (Fig. 1B). Urine concentration, in part, depends on the uptake of water through AQP2. Trafficking and regulation of AQP2 into the membrane occurs through the activation of the vasopressin receptor or hyperosmotic increase [38]. Upon activation, vesicles travel through the microtubules and are inserted in the apical membrane [39]. So far, we have shown that deletion of PC2 disrupts the tethering of the microtubule to the ER and impairs microtubules dynamics (Fig. 4). To determine the physiological effect of the PC2-MAP4 osmosensing pathways, we examined AQP2 distribution after hyperosmotic challenge. At baseline in CTL cells, AQP2 resided in the cytosol with minimal labeling in the membrane (Fig. 7A, *0 min*). After stimulating the cells for 10 min with hyperosmotic solution, AQP2 puncta moved to the apical membrane (Fig. 7A, *10 min*). Additionally, hyperosmotic stimuli caused MAP4 co-localization with AQP2 in the membrane of CTL cells. In contrast, there was minimal labeling of AQP2 and MAP4 in the membrane of PC2 KO cells, and this was unchanged during the hyperosmotic challenge (Fig. 7B). Increase of extracellular osmolarity significantly increased AQP2 cytosolic vesicle size in imCD3 CTL cells 10 min after exposure (Fig. 7C). In contrast, AQP2 cytosolic vesicles were already increased at baseline in imCD3 PC2 KO cells and remained unchanged after osmotic stimuli (Fig. 7C). As a control, we stimulated the cells with AVP (100nM) to activate the canonical pathway of AQP2 trafficking into the membrane. In CTL cells, both AQP2 and MAP4 co-labeled in the membrane after 10 min with AVP (Fig. 7D). Interestingly, in PC2 KO cells, half of the cells showed labeling of AQP2 in the membrane, but not of MAP4 (Fig. 7D). We confirmed these results by examining the localization of AQP2 in the collecting ducts of CTL and PC2 KO mice (Fig. 7E-F). Labeling of AQP2 and MAP4 in CTL mice localized in the membrane (Fig. 7E). In comparison, to the tubule PC2 KO mice, there was diffuse cytosolic staining of AQP2 and MAP4 (Fig. 7F). In conclusion, we observed that the tethering of the cytoskeleton and the ER mediated by PC2 and MAP4 have biological relevance as the disruption of these interactions impaired the trafficking of AQP2 channel into the membrane, therefore impacting the ability to concentrate urine.

**Figure 7.**
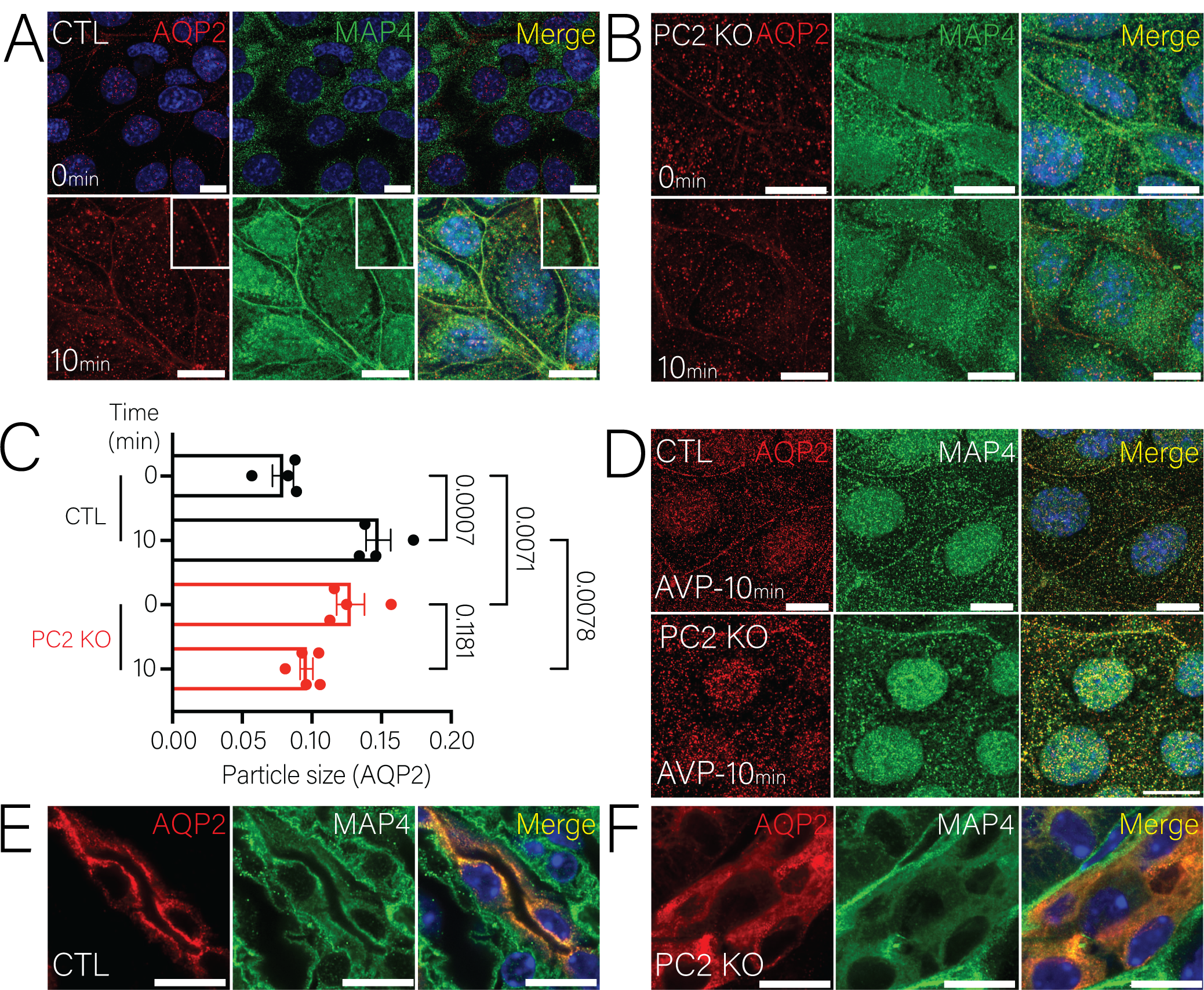
Aquaporin 2 trafficking into the membrane is impaired in PC2 KO cells and in collecting ducts from PC2 KO mice. **A.** Representative immunofluorescent staining of imCD3 CTL cells at basal osmotic conditions (top panels) labeling AQP2 (red) and MAP4 (green) and 10 min after increasing extracellular osmolarity to 400 mOsm (bottom panels). Scale bars represent 10 μm. **B.** Immunofluorescent staining of A (red) and MAP4 (green) in PC2 KO imCD3 cells at basal osmotic conditions (top panels) and after increase of extracellular osmolarity (bottom panels). Scale bars represent 10 μm. **C.** Quantification of AQP2 vesicle size significantly increases in imCD3 CTL cells after increasing extracellular osmolarity (400 mOsm) for 10 min. Vesicle size remained unchanged in PC2 KO cells. Bars represent mean±SEM. Data were analyzed to determine normality and statistical analysis was determined by 2-way ANOVA. p-values listed in the figure. **D.** Immunofluorescent staining of AQP2 insertion in the presence of 100nM AVP of PC2 KO imCD3 cells. Scale bars represent 10 μm. **E.** Immunofluorescent staining of AQP2 (red) and MAP4 (green) in isolated renal tubules from CTL mice. AQP2 and MAP4 colocalized into the membrane of the collecting duct cell membrane (top panels). Scale bars represent 10 μm. **F.** Immunofluorescent staining of AQP2 (red) and MAP4 (green) in isolated renal tubules from tubule specific PC2 KO mice. AQP2 and MAP4 have diffused staining in the cytosol of collecting duct cells (top panels). Scale bars represent 10 μm.

## Discussion

The physiological function of the polycystin TRP channel, PC2, has been contentious. Although the clinical importance of PC2 is well established, as mutations to PC2 result in the genetic disorder ADPKD, the functional role and the identification of a specific agonist that activates PC2 has been unclear. Many studies have established that PC2 localizes to primary cilium where it appears to mediate ciliary signaling by itself or in complexes with PC1 or with other TRP channels like TRPM3 [10, 17]. However, whether ciliary PC2 conducts calcium under physiological conditions has been debated. More importantly, a bigger question that remains unanswered is the conditions that lead to the activation of PC2 channel activity, either in the cilia, or the ER, where its role remains poorly understood. Our study provides a model that provides answers to both questions: what is a physiological stimulus that activates PC2, and what is the role of PC2 within the ER. We demonstrate that hyperosmotic shifts activate calcium release through PC2 channels in the ER by a novel mechano-transduction pathway that links the microtubules to the ER through the interaction of PC2 and MAP4. Our data demonstrates that hyperosmotic stimuli induce microtubule depolymerization which in turn “tugs” on PC2 allowing for calcium release [40]. The initial calcium release by PC2 recruits other ER localized calcium channels, most likely the InsP_3_R, which sustain the cytosolic calcium response. Functionally, this osmosensing pathway allows for the insertion of AQP2 in the membrane of collecting duct cells in an AVP-independent manner, which is of high physiologic relevance as pre-cystic PC2 KO mice, like humans with ADPKD, have decreased urine concentrating ability.

PC2 and its partner protein PC1 have been described as mechanosensory proteins, in part due to its localization in the primary cilia [19], which can bend in response to changes in fluid flow or sheer stress. It has been thought that this bending can open the PC2 channel, or the PC1/PC2 complex in cilia, and presumably flux calcium, but as a non-selective ion channel, it is possible that monovalent ions like sodium can be conducted. Although electrophysiological approaches have tried to discern the ionic preference of PC2 in the primary cilia, the conditions required to activate PC2 do not resemble physiological conditions that the cell will experience [14–16, 18]. Therefore, the activation of the channel activity of PC2 to determine its ionic preference, although important, has become disconnected from the physiological conditions that could potentially activate PC2 and provide insight into the physiological roles these proteins play in renal cells. We therefore chose to use a mild hyperosmotic stimulus (400 mOsm), which represents a condition that normal collecting duct epithelial cells would be expected to experience. Surprisingly, although the collecting duct can experience a wide range in osmolarities (between 50-1200 mOsm), there have been no comprehensive studies as to whether polycystin proteins are activated by osmotic shifts. Although prior literature has demonstrated that osmotic shifts within renal epithelial cells cause a rise in intracellular calcium, this has not been connected to the polycystin proteins [41]. We show that isolated kidney sections and renal epithelial cells (and a variety of different cell lines) with PC2 deletions, were unable to increase their cytosolic calcium in response to extracellular osmotic shifts. Importantly, for the wider discussion on the function of PC2 in the cilia, we found that hyperosmotic stimuli by itself was incapable of eliciting a ciliary calcium increase (Fig. 3). Moreover, the increase of cytosolic calcium that we did see by the hyperosmotic stimuli does not back-propagate to the cilia (which has been a scenario proposed by other investigators). The very tight regulation of the calcium response to hyperosmotic shifts highlights the distinction of ER-localized PC2 versus ciliary-localized PC2. It is possible that ciliary PC2 is required to sense other environmental cues such as changing fluid flow, where repeated bending of the cilia might allow for calcium influx through PC2 [20, 21]. Thus, our proposed model is in harmony with the ciliary hypothesis as it appears that the two localizations of PC2-on the cilia, and on the ER-can discriminate between the extracellular cues that the cells are experiencing (Fig 8). Therefore, PC2 in the ER can respond to changes of osmolarity, whereas PC2 in the cilia can be activated by multiple bending of the cilia. This would be congruent with previous data where 30 bends are required to induce a nodal ciliary calcium response via PC2 [20, 21]. However, we cannot discount the fact that osmotic changes can activate signaling pathways within the cilia that are calcium independent, like Wnt, Notch and Hedgehog signaling [42]. Therefore, the fact that ciliary calcium does not increase upon osmotic changes, does not necessarily indicate that other signaling pathways are being activated. Future studies would require the combination of both fluid flow and osmotic changes to determine whether PC2 in the cilia acts as a coincident detector of osmolarity and fluid flow.

Another question that arises from our findings is the role of TRP channels that are activated by osmolarity and are present in renal cells. Hypoosmotic stimulus has been shown to lead to calcium influx through both TRPC3 and TRPV4 [41, 43, 44]. It is possible that hypoosmotic stimuli (which presumably would have a very different mechanistic pathway of activation than the hyperosmotic stimuli tested here) could elicit activation of a different set of TRP channels. However, recent work proposed that hyperosmotic changes can also lead to the activation of calcium influx via TRPV4 [43]. Nonetheless, the activation of calcium influx via hyperosmotic stimuli was not comparable to the calcium influx activated by TRPV4 specific agonists [43]. More importantly, the authors suggested that the cytosolic calcium increase induced by hyperosmolarity can be due to other calcium channels. As PC2 has been shown to heterodimerize with TRPV4 [45] it is possible that the requirement for TRPV4 in hyperosmotic stimuli also necessitates PC2.

Our results also reveal a novel mechanosensation pathway whereby the hyperosmotic stimuli reduce tension on the microtubules. Prior studies in the literature have suggested that hyperosmotic changes induce fast changes in membrane tension (order of seconds to minutes) followed by a volumetric change which is a separate and longer process (on the order hours) [46]. The speed of the response that we characterize in this study is driven by changes in membrane tension which demonstrate that the organization of the microtubules and dynamics are significantly impaired and decreased in PC2 KO cells (Fig. 3). This could suggest that the microtubules in the PC2 KO cells are unable to sense changes in tension which can transmit the message to the plasma membrane opening additional channels to allow for calcium influx. Therefore, although other TRP channels may be involved in mediating hypoosmotically induced calcium signaling, our data demonstrate that ER-localized PC2 in concert with the microtubules are responsible for mediating the intracellular calcium release in response to hyperosmolarity. It remains to be determined if ER-localized PC2 also contributes to hypoosmotically driven calcium responses.

Osmotic changes induce the activation of cellular processes that are required for cell adaptation, not just for renal epithelial cells but cells in other environments [47]. Previous work on the PC2 homologue in fission yeast, Pkd2, was previously shown to be activated via osmotic stimuli [48]. Additionally, it has been shown that interaction of PC2 with cytoskeletal components can regulate PC2 channel activity [49–52]. Why is the activation of ER-localized PC2 by hyperosmotic stimuli in renal epithelial cells important? In ADPKD patients, one of the early symptoms experienced is decreased urine concentrating abilities [53]. Urine concentration in part occurs by the uptake of water into the cells through the incorporation of AQP2 into the membrane through canonical (AVP-dependent) and non-canonical pathways (like hyperosmolarity) [54]. Previous work has pointed to the higher expression of AQP2 in ADPKD models, which were not based on mutations or loss of polycystin proteins [55, 56]. Therefore, how this pathway is affected in ADPKD model remains unknown. In this study, we provide a mechanism by which the tethering of the microtubules to the ER, mediated by PC2 and MAP4, is disrupted by phosphorylation of MAP4 with hyperosmotic stimulus, enabling AQP2 trafficking to the membrane through a non-AVP pathway and that this mechanism of AQP2 trafficking is impaired in a pre-cystic PC2 KO ADPKD model. Previous work has demonstrated that in addition to stabilizing microtubules, MAP4 regulates the directionality of the cargo being transported through the microtubules [57–59]. We demonstrate that there is a reciprocal dependence in MAP4/PC2 expression. When PC2 is knocked out, MAP4 expression is decreased, whereas deletion of MAP4 increases PC2 expression (Supp. Fig. 9). Therefore, both phosphorylation and decreased MAP4 expression not only affects stability of the microtubules but the ability of cargo, like AQP2 vesicles, to be transported. Moreover, the transport machinery that allows for these vesicles to travel through the microtubules are calcium dependent [60]. Therefore, the absence of calcium signals in the PC2 KO cells could additionally impair the transport machinery further exacerbating the urine concentration deficiency as AQP2 channels are unable to be incorporated into the membrane. Collectively, loss of these functions would lead to dilute urine, which is seen in both our study here and in a pre-cystic PC1 KO mouse [61] as well as ADPKD patients.

In this study we provide a physiological role of ER-localized PC2 in renal cells. We demonstrate that the interaction between the ER and microtubules, mediated by PC2 and MAP4, are critical for signaling pathways like the incorporation of AQP2 into the membrane of renal cells (Fig. 8). The novel interaction between PC2 and MAP4 provides not only structural support for organelles, but the mechanism by which these external cues are transmitted to intracellular signals. Here we demonstrate the physiological implications of these interactions by linking how deletion of PC2 leads to decreased urine osmolarity. These findings may also provide a contributing mechanism by which circulating vasopressin levels are higher in ADPKD patients, as the collecting duct cells have impaired water uptake, which in turn, will drive higher levels of circulating vasopressin. Indeed, the only approved treatment for ADPKD that halts cyst growth is tolvaptan which targets the V2R [62, 63], and for which AQP2 is the downstream target. Thus, our findings provide foundational insights into PC2 biology by which impairment of urine concentration can contribute to cyst development. In conclusion, hyperosmolarity is a natural agonist that activates calcium release from ER-localized PC2 in the osmosensing pathway by its interaction with the microtubules.

## Acknowledgements

We thank Dr. Erika Piedras Rentería, Dr. Jordan Beach, Dr. Patrick Oakes, Dr. Barbara Ehrlich, Dr. Arlene Chapman and Kuo lab members for helpful discussions. Reagents: Dr. David Yule (University of Rochester, HEK 293 TKO cells), Dr. Aldebaran Hofer (Harvard Medical School, gCaMP6F-Arl13B), Dr. Jordan Beach (Loyola University Chicago, gCaMP7s-CAAX), Dr. Aleksey Zima (Loyola University Chicago, R-cepia) and Dr. Stefan Somlo (Yale University, Pkd2 floxed mice). We thank the Department of Cell and Molecular Physiology (Loyola University Chicago) for access to the specialized imaging resource center (Zeiss 880-Airyscan) and metabolic phenotypic resource center (metabolic cages). We thank Dr. Charlie Yang (Rosalind Franklin University) for conducting the mass spectrometry analysis. Grants and funding: R00DK101585 (IYK); Research reported in this publication was supported by the National Institute of Diabetes and Digestive and Kidney Diseases of the National Institutes of Health under Award Numbers U2CDK129917 and TL1DK132769. (KMMN).

## Contributions

Conception: KMMN and IYK. Experiments: KMMN, RMK, VV, IYK. Analysis: KMMN, IYK. Discussions: KMMN, RMK, VV, IYK. Draft: KMM and IYK. All authors approved the final manuscript.

---

**Supplementary Figure 1.**
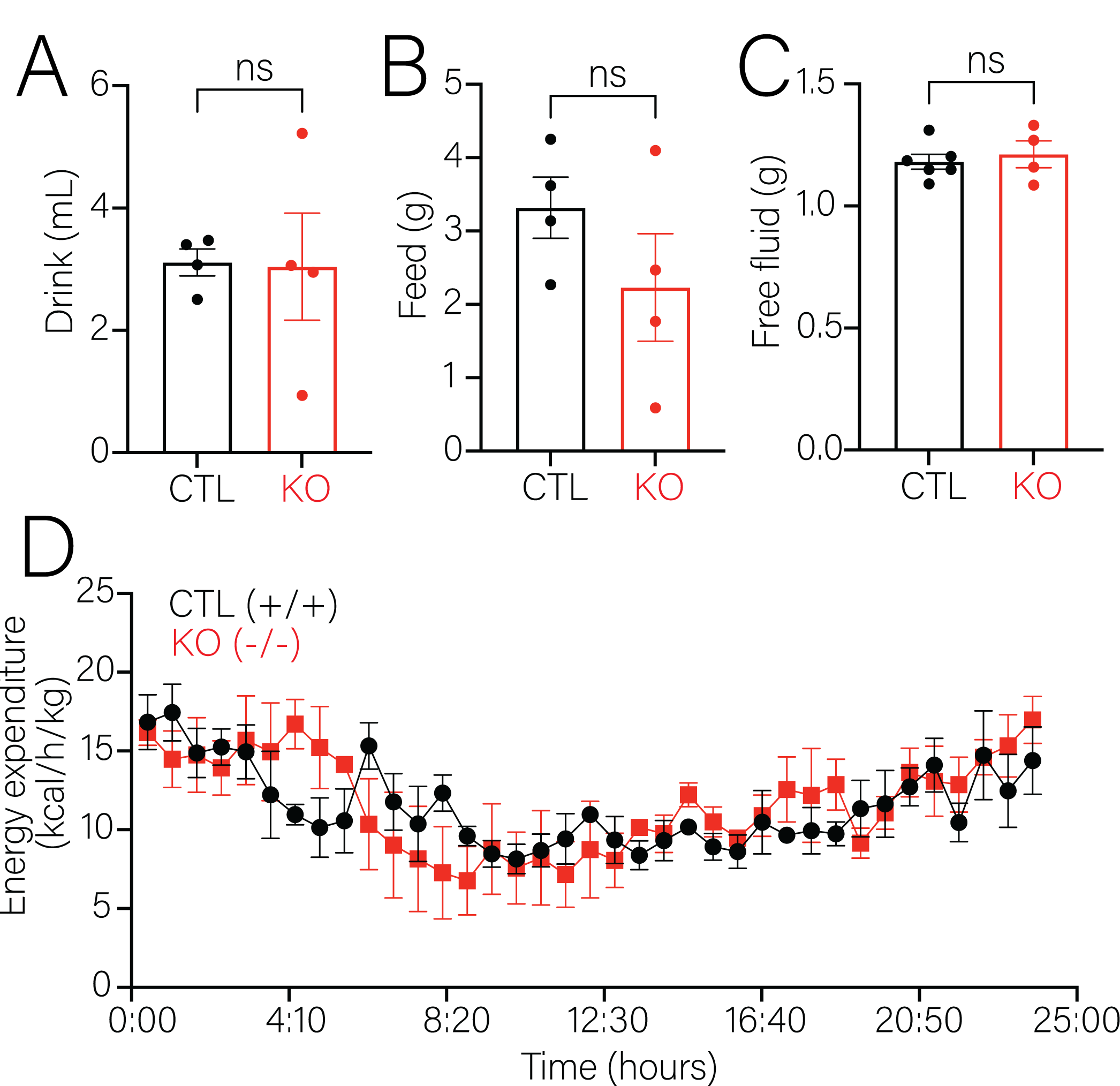
Water consumption in CTL and tubule specific PC2 KO mice. **A.** Consumption of water during 24 hours of CTL and tubule specific PC2 KO mice in metabolic cages. Bars represent mean±SEM. Data were analyzed to determine normality. Statistical analysis was determined by student’s t-test. **B.** Food consumption during 24 hours of CTL and tubule specific PC2 KO mice in metabolic cages. Bars represent mean±SEM. Data were analyzed to determine normality. Statistical analysis was determined by student’s t-test. **C.** Free bodily fluid measured through NMR 24 hours before introduction to metabolic cages of CTL and tubule specific PC2 KO mice. Bars represent mean±SEM. Data were analyzed to determine normality. Statistical analysis was determined by student’s t-test. **D.** Analysis of energy expenditure during a 24-hour period of CTL and tubule specific PC2 KO mice in metabolic cages. Bars represent mean±SEM. Data were analyzed to determine normality. Statistical analysis was determined by student’s t-test.

**Supplementary Figure 2.**
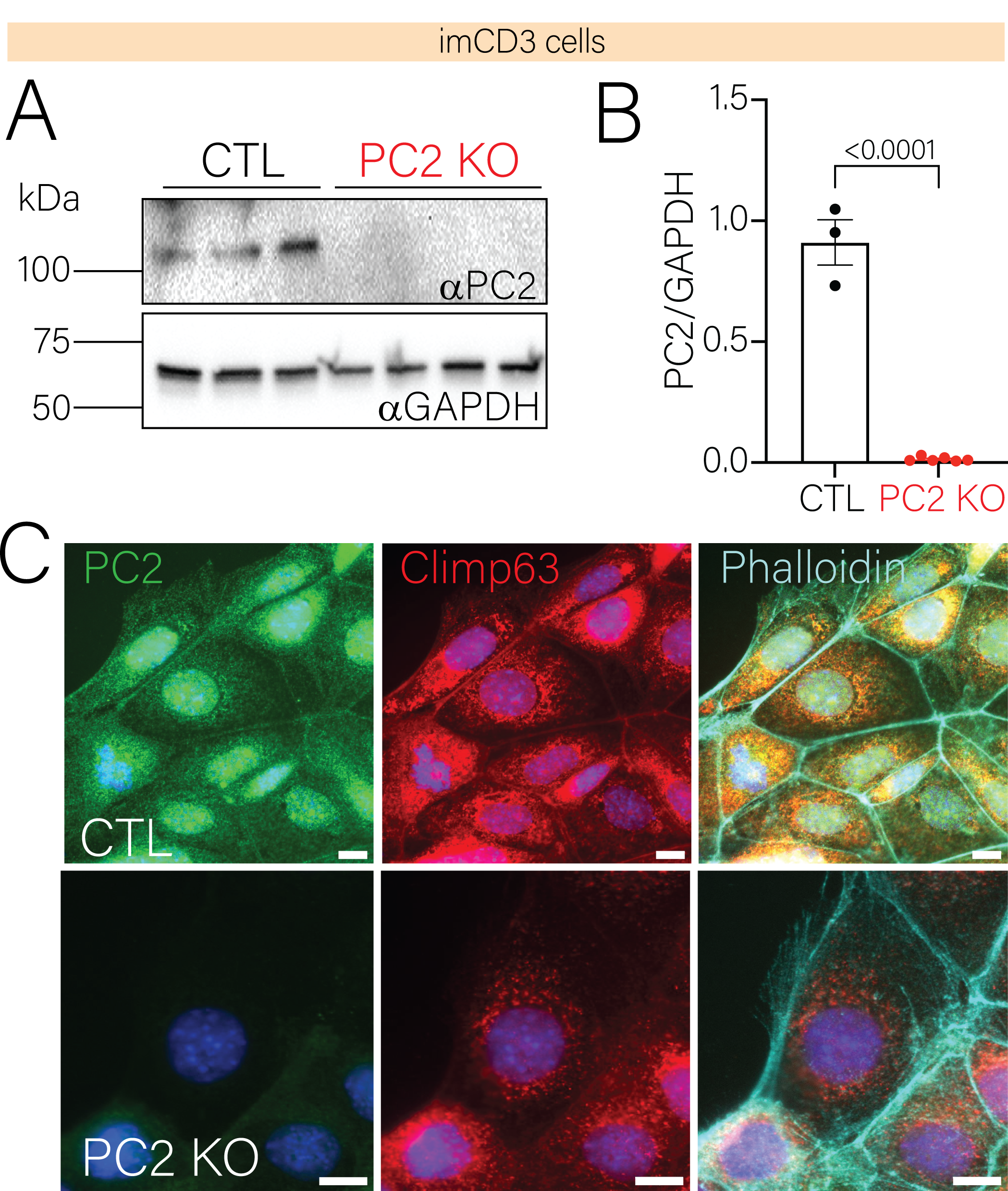
Validation of PC2 KO in imCD3 cells. **A.**Expression of PC2 in imCD3 CTL and PC2 KO cells. GAPDH was used as loading control. **B.** PC2 expression was significantly decreased in PC2 KO imCD3 cells. **C.** Immunofluorescent staining of PC2 (green) and CLIMP63 (ER morphology; red) and phalloidin (cyan) in imCD3 CTL (top panel) and PC2 KO cells (bottom panels). Scale bars represent 10 μm.

**Supplementary Figure 3.**
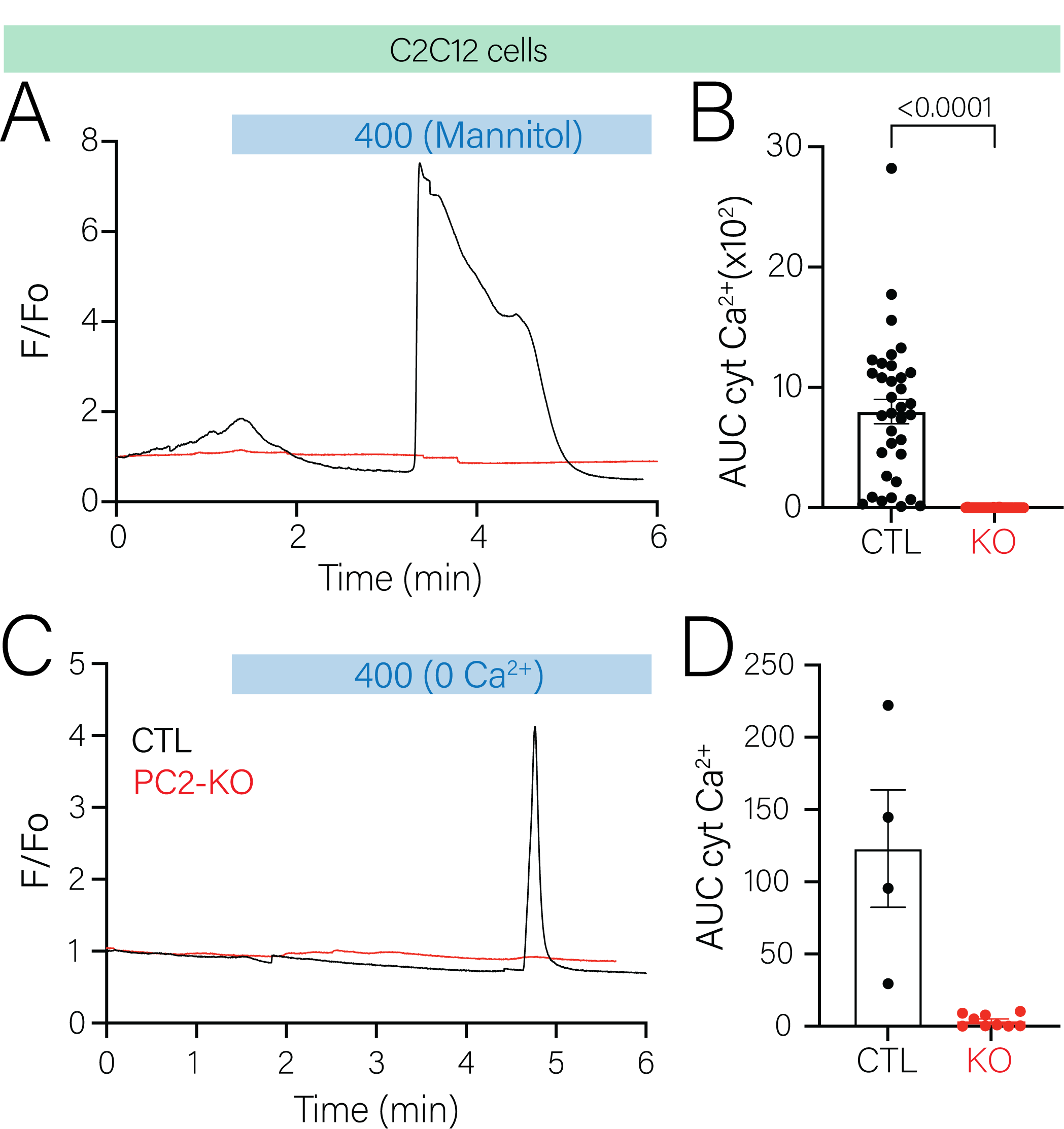
Hyperosmotic stimuli induced intracellular calcium release. **A.** Representative trace of cytosolic calcium changes in CTL (*black line*) and PC2 KO C2C12 cells (*red line*) after increase of extracellular osmolarity using mannitol. Cytosolic calcium increase was absent in PC2 KO cells. **B.** Area under the curve was significantly decreased in C2C12 PC2 KO cells. Bars represent mean±SEM. Data were analyzed to determine normality. Statistical analysis was determined by student’s t-test. p-values listed in figure. **C.** Representative trace of cytosolic calcium changes in CTL (*black line*) and PC2 KO C2C12 cells (*red line*) after increase of extracellular osmolarity in the absence of extracellular calcium. **D.** Area under the curve was significantly decreased in C2C12 PC2 KO cells. Bars represent mean±SEM. Data were analyzed to determine normality. Statistical analysis was determined by student’s t-test. p-values listed in figure.

**Supplementary Figure 4.**
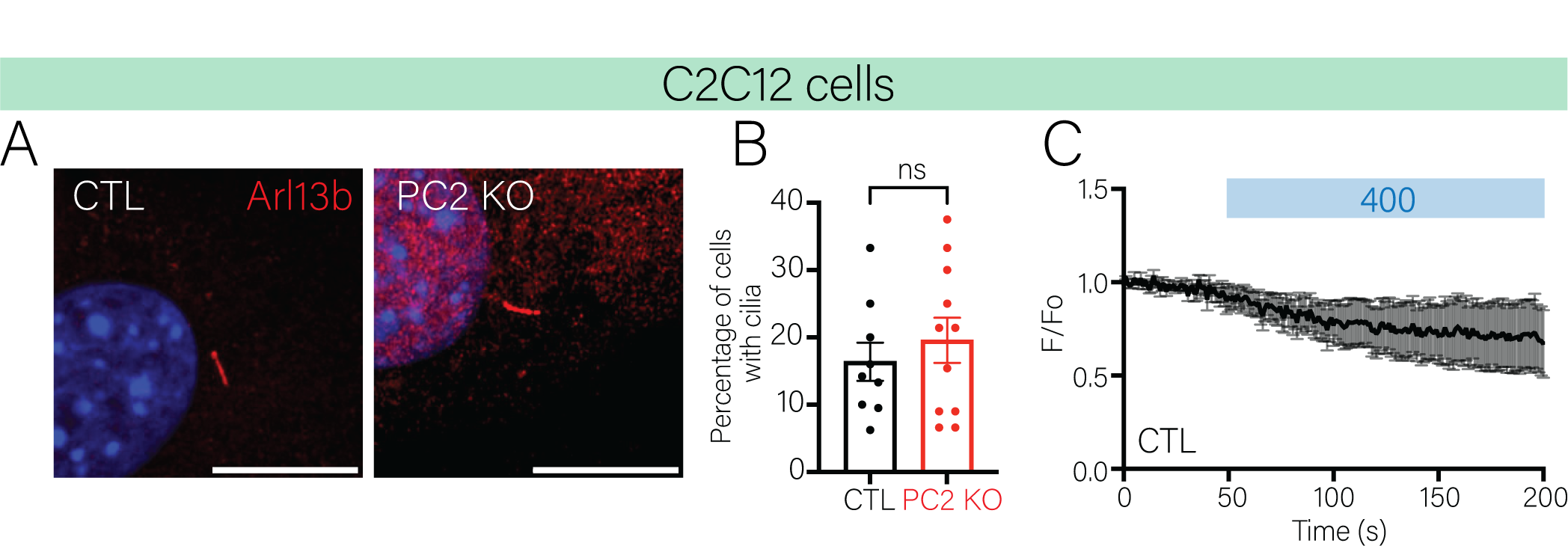
Deletion of PC2 does not affect cilia formation. **A.** Representative immunofluorescent staining of Arl13B in C2C12 CTL (left panel) and PC2 KO cells (right panel). Cells were stained 24 hours after plating mimicking the same conditions as the cytosolic calcium assay. Scale bars represent 10 μm. **B.** Quantification of the percentage of cilia per total number of cells was similar between C2C12 CTL and PC2 KO cells. C. Ciliary calcium did not increase in CTL C2C12 cells after increase of extracellular osmolarity to 400 mOsm.

**Supplementary Figure 5.**
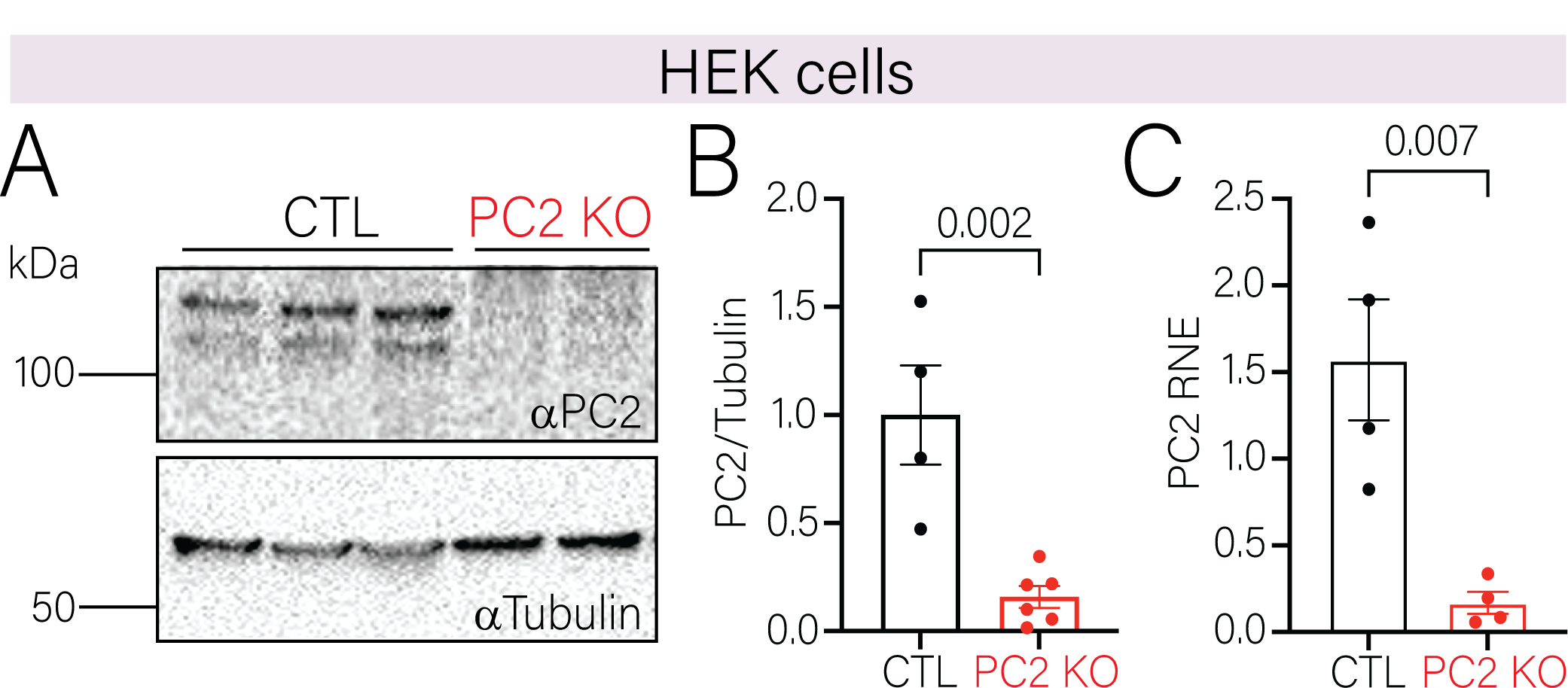
Validation of PC2 KO in Human embryonic kidney (HEK) cells. **A.** Expression of PC2 in CTL and PC2 KO cells in HEK cells. Tubulin was used as loading control. **B.** PC2 expression was significantly decreased in PC2 HEK cells. Bars represent mean±SEM. Data were analyzed to determine normality. Statistical analysis was determined by student’s t-test. p-values listed in the figure. **C.** mRNA of PC2 was significantly decreased in PC2 KO HEK cells. Bars represent mean±SEM. Data were analyzed to determine normality. Statistical analysis was determined by student’s t-test. p-values listed in the figure.

**Supplementary Figure 6.**
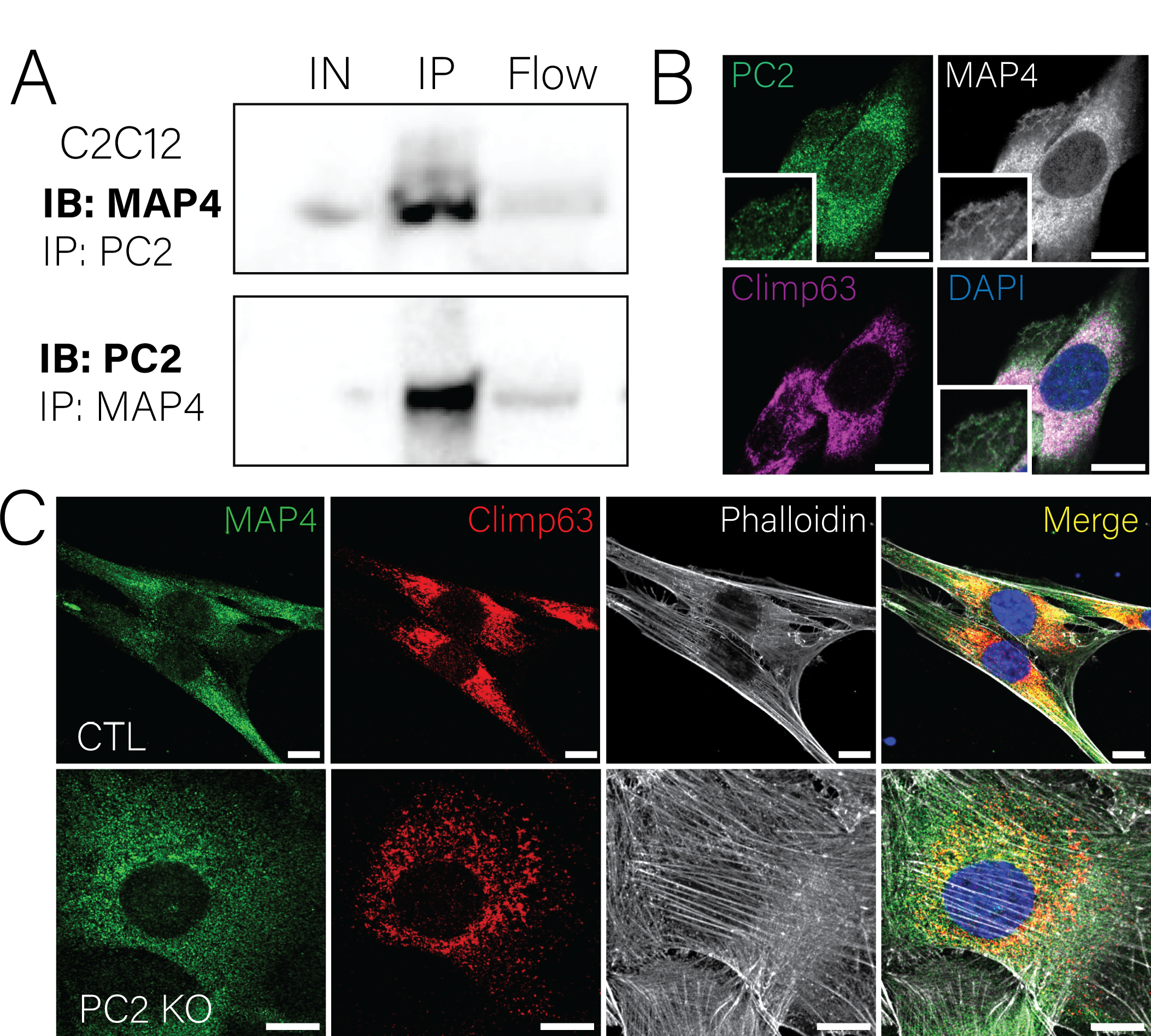
PC2 interacts with microtubule binding protein 4 (MAP4) **A.** Reciprocal immunoprecipitation assay validating interaction between PC2 and MAP4. IN: input; IP: immunoprecipitation. **B.** Immunofluorescent staining of PC2 (green), MAP4 (white) and CLIMP63 (ER; magenta) in CTL C2C12 cells. **C.** Immunofluorescent staining of MAP4 (green), CLIMP63 (ER; red) and phalloidin (white) in C2C12 CTL cells (top panels) and PC2 KO cells (bottom panels). Scale bars represent 10 μm.

**Supplementary Figure 7.**
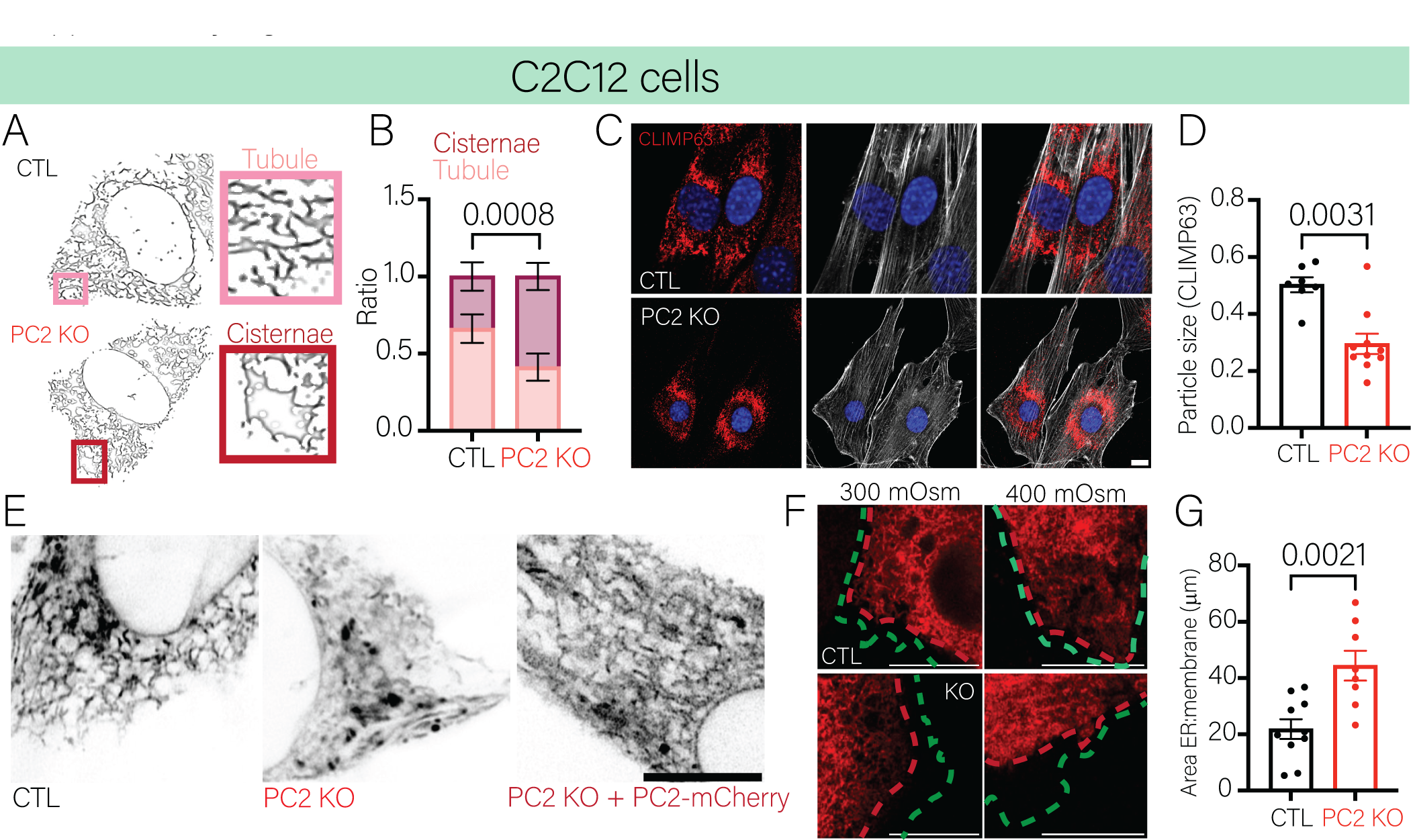
Deletion of PC2 affects ER morphology. **A.** Processed high resolution images of C2C12 CTL and PC2 KO cells expressing ER-mCherry. Inset: Tubular morphology (tubules) and cisternaes (falsely colored). **B.** The ratio between tubules and cisternae was significantly increased in C2C12 PC2 KO cells. Bars represent mean±SEM. Data were analyzed to determine normality. Statistical analysis was determined by Two-way ANOVA followed by Sidak’s. p-values listed in figure. **C.** Immunofluorescent staining of C2C12 CTL (top panels) and PC2 KO cells (bottom panels) of CLIMP63 (ER, red) and phalloidin. **D.** CLIMP63 particle size was significantly decreased in C2C12 PC2 KO cells in comparison to the CTL cells. Bars represent mean±SEM. Data were analyzed to determine normality. Statistical analysis was determined by Mann-Whitney’s test (Gaussian distribution was not assumed. p-values listed in figure. **E.** ER morphology of C2C12 CTL and PC2 KO cells using ER-mCherry. Re-expression of full-length PC2 (PC2-mCherry) restored tubular morphology in PC2 KO cells. **F.** Representative images of C2C12 CTL Cells (top panels) and PC2 KO cells (bottom panels) at 300 mOsm (left) and 400 mOsm (right) labeling the ER (ER-mCherry, red) and the plasma membrane (gCaMP7s-CAAX, green). Dotted lines represent the border of the ER membrane or plasma membrane. Scale bars represent 10 μm. **G.** Quantification of the area between the plasma membrane and the ER were significantly increased in the PC2 KO cells. Bars represent mean±SEM. Data were analyzed to determine normality. Statistical analysis was determined by student’s t-test. p-values listed in figure.

**Supplementary Figure 8.**
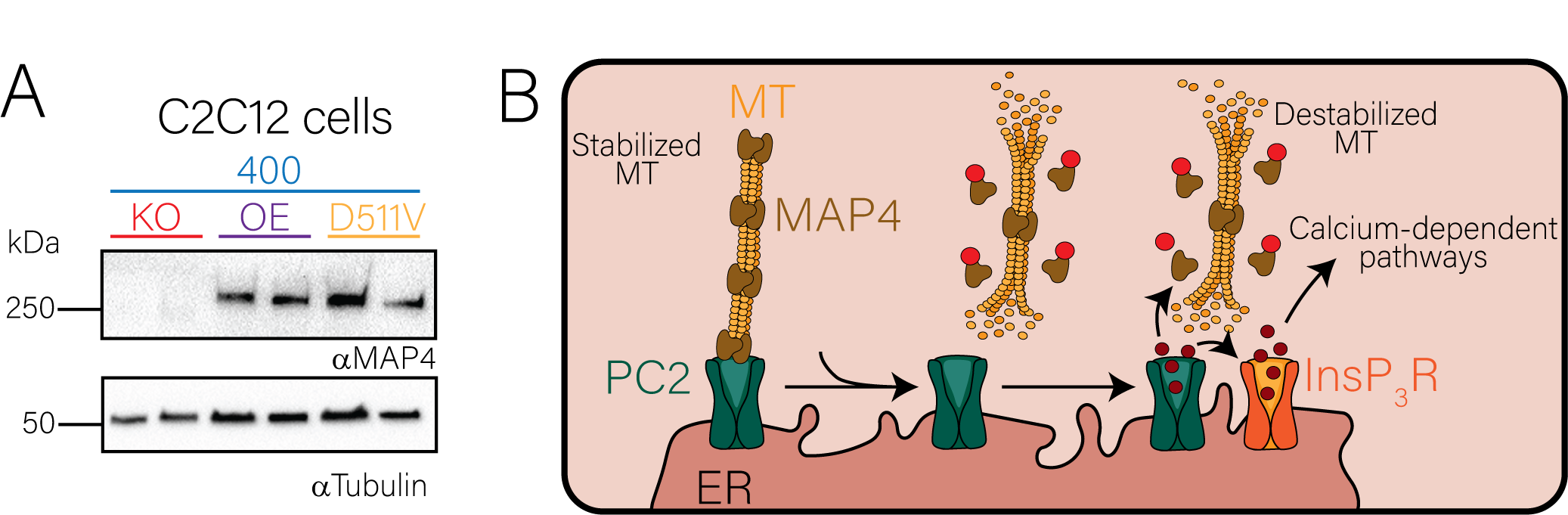
MAP4 expression was restored upon expression of full-length PC2. **A.** Expression of MAP4 and CLIMP63 in C2C12 PC2 KO cells, PC2 KO + PC2 (OE, purple) and PC2 KO + PC2 D511V (D511V/D; orange) at 400 mOsm. GAPDH was used as loading control. **B.** Model of how osmolarity induces MAP4 disassociation from the microtubules which destabilizes them. Destabilization then “tugs” PC2 which opens allowing for calcium release and recruiting InsP3R to sustain the cytosolic calcium increase.

**Supplementary Figure 9.**
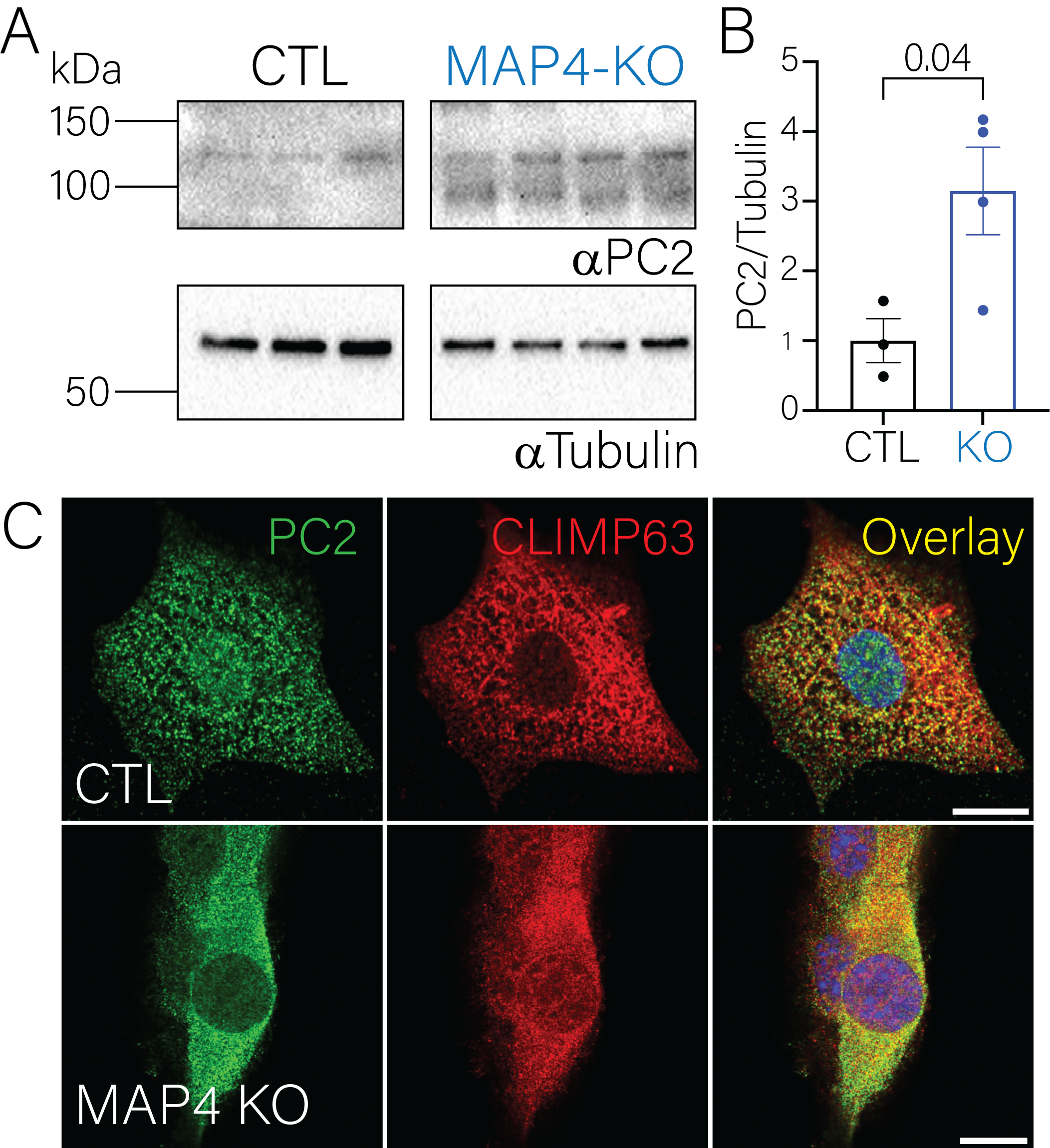
PC2 expression increases upon knock out of MAP4. **A.** Expression of PC2 in C2C12 CTL and MAP4-KO cells. Tubulin was used as loading control. **B.** PC2 expression was significantly increased in MAP4-KO cells in comparison to CTL cells. Bars represent mean±SEM. Data were analyzed to determine normality. Statistical analysis was determined by student’s t-test. p-values listed in figure. **C.** Immunofluorescent staining of PC2 (green) and CLIMP63 (ER; red) in C2C12 CTL cells (top panels) and MAP4-KO cells (bottom panels). Scale bars represent 10 μm.

**Supplementary Table 1.** Protein list of PC2 immunoprecipitation mass spectrometry analysis.

| Accession | Coverage (%) | Peptides | Avg. Mass | Description |
| --- | --- | --- | --- | --- |
| Q62261\|SPTB2_MOUSE | 19 | 39 | 274221 | Spectrin $\beta$ chain non-erythrocytic 1 |
| P15508\|SPTB1_MOUSE | 15 | 25 | 245248 | Spectrin $\beta$ chain erythrocytic |
| A2AQP0\|MYH7B_MOUSE | 6 | 34 | 221495 | Myosin-7B |
| P16546\|SPTN1_MOUSE | 11 | 22 | 284596 | Spectrin $\alpha$ chain non-erythrocytic 1 |
| P70670\|NACAM_MOUSE | 13 | 19 | 220497 | Nascent polypeptide-associated complex subunit alpha muscle-specific form |
| P27546\|MAP4_MOUSE | 25 | 15 | 117429 | Microtubule-associated protein 4 |
| P02468\|LAMC1_MOUSE | 9 | 11 | 177298 | Laminin subunit gamma 1 |
| O55143\|AT2A2_MOUSE | 7 | 6 | 114858 | Sarcoplasmic/endoplasmic reticulum calcium ATPase 2 |

**Supplementary Video 1:** Osmotic calcium induced rise in collecting ducts from renal tubules of CTL and PC2 KO mice.

**Supplementary Video 2:** Osmotic calcium induced rise in CTL and PC2 KO immortalized murine collecting duct cells (imCD3).

**Supplementary Video 3:** Hyperosmotic stimuli shortens EB3 comets in C2C12 CTL and PC2 KO cells.

